## Supplementary Figures and Tables for "Oviductin sets the species-specificity of the mammalian zona pellucida"

Daniel de la Fuente *et al*.

**This PDF file includes:**

Figs. S1 to S8

Tables S1 to S14

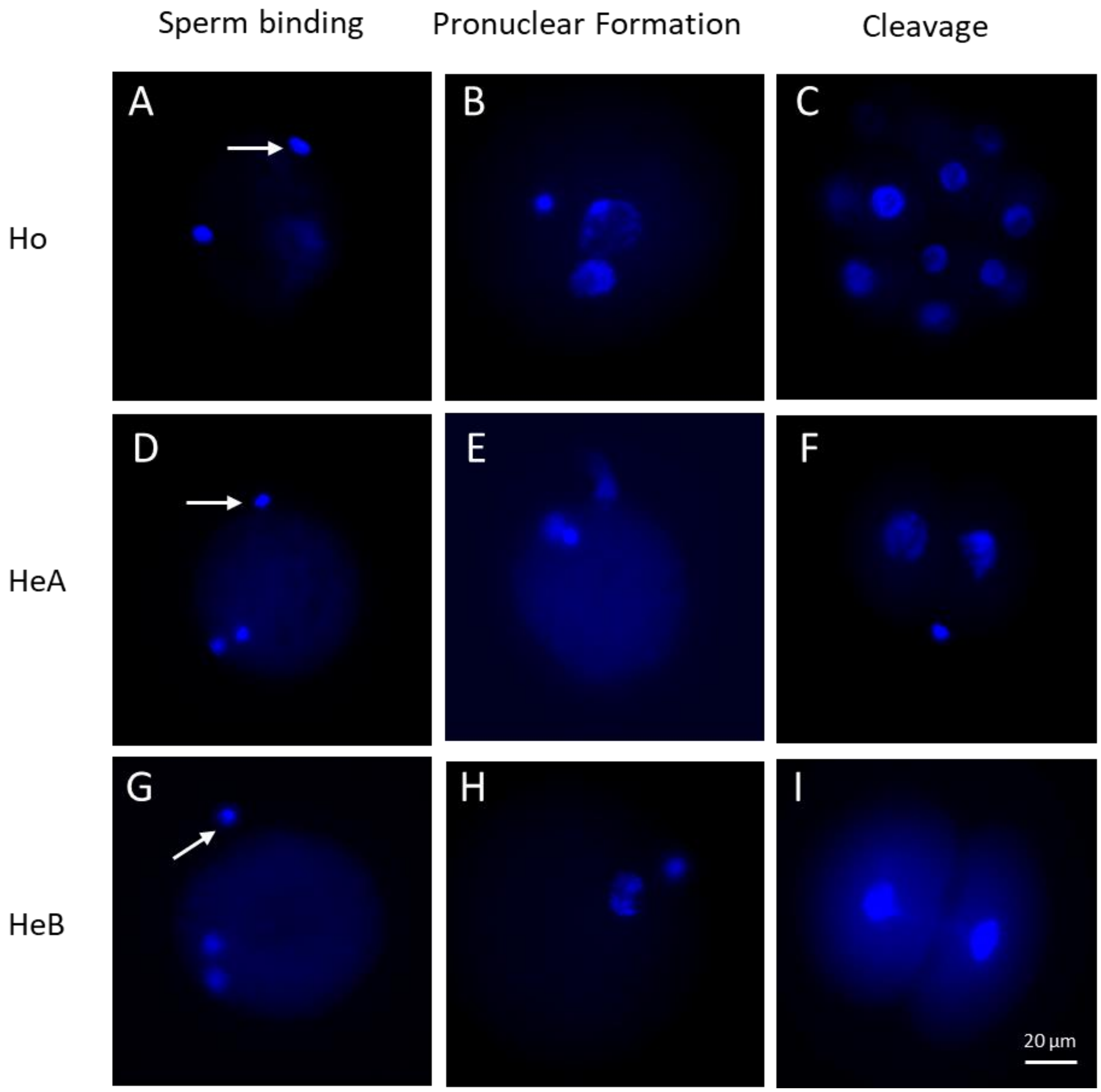


**Fig. S1. Heterologous IVF between bovine oocytes and human sperm.** Sperm-oocyte binding, pronuclear formation, and cleavage after homologous (Ho) (bovine sperm) and heterologous (He) IVF: human sperm capacitated with G-IVF™ PLUS medium (HeA) or Tyrode's medium (HeB). Gametes were stained with Hoechst 33342. Ho: Bound bovine sperm after 2.5 hours of co-incubation with zona-intact bovine oocytes (**A**); pronuclear formation at 6 hpi (**B**); and embryo cleavage at 48 hpi (**C**). HeA and HeB: Bound human sperm after 2.5 hours of co-incubation of the gametes (**D**, **G**); pronuclear formation at 18 hpi (**E**, **H**); and hybrid-embryo cleavage at 48 hpi (**F**, **I**), respectively. Arrow points to sperm head chromatin. Images were captured with a 63X objective. Scale bar 20 µm.

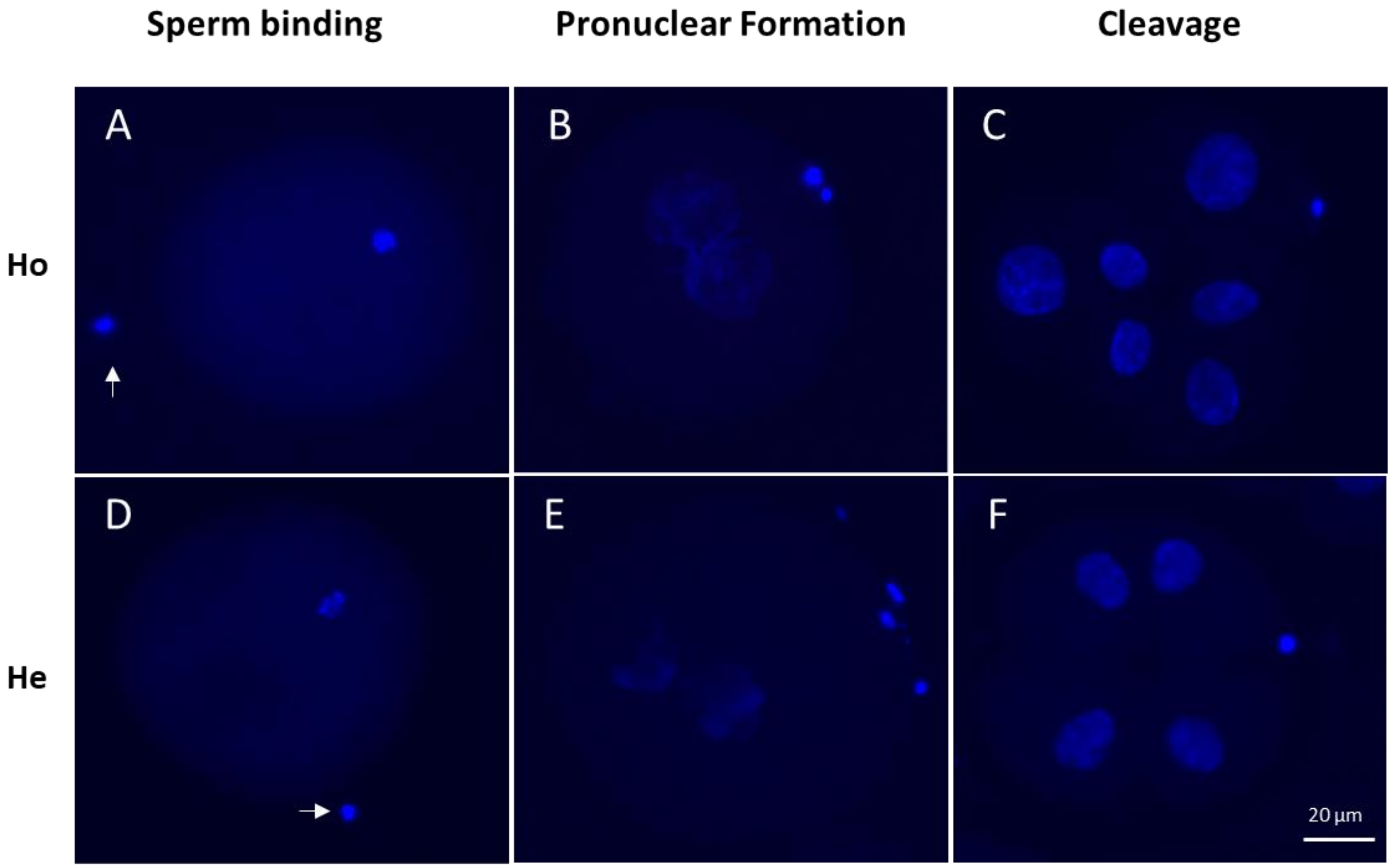


**Fig. S2. Heterologous IVF between bovine oocytes and murine sperm.** Sperm-oocyte binding, pronuclear formation, and cleavage after homologous (Ho, bovine sperm) and heterologous (He, mouse/rodent sperm) IVF. Gametes were stained with Hoechst 33342. Ho: Bound bovine sperm after 2.5 hours of co-incubation with zona-intact bovine oocytes (A); pronuclear formation at 6 hpi (B); and embryo cleavage at 24 hpi (C). He: Bound mouse sperm after 2.5 hours of co-incubation of the gametes (D); pronuclear formation at 18 hpi (E); and hybrid-embryo cleavage at 24 hpi (F). Arrow points to sperm head chromatin. Images were captured with a 63X objective. Scale bar 20 µm.

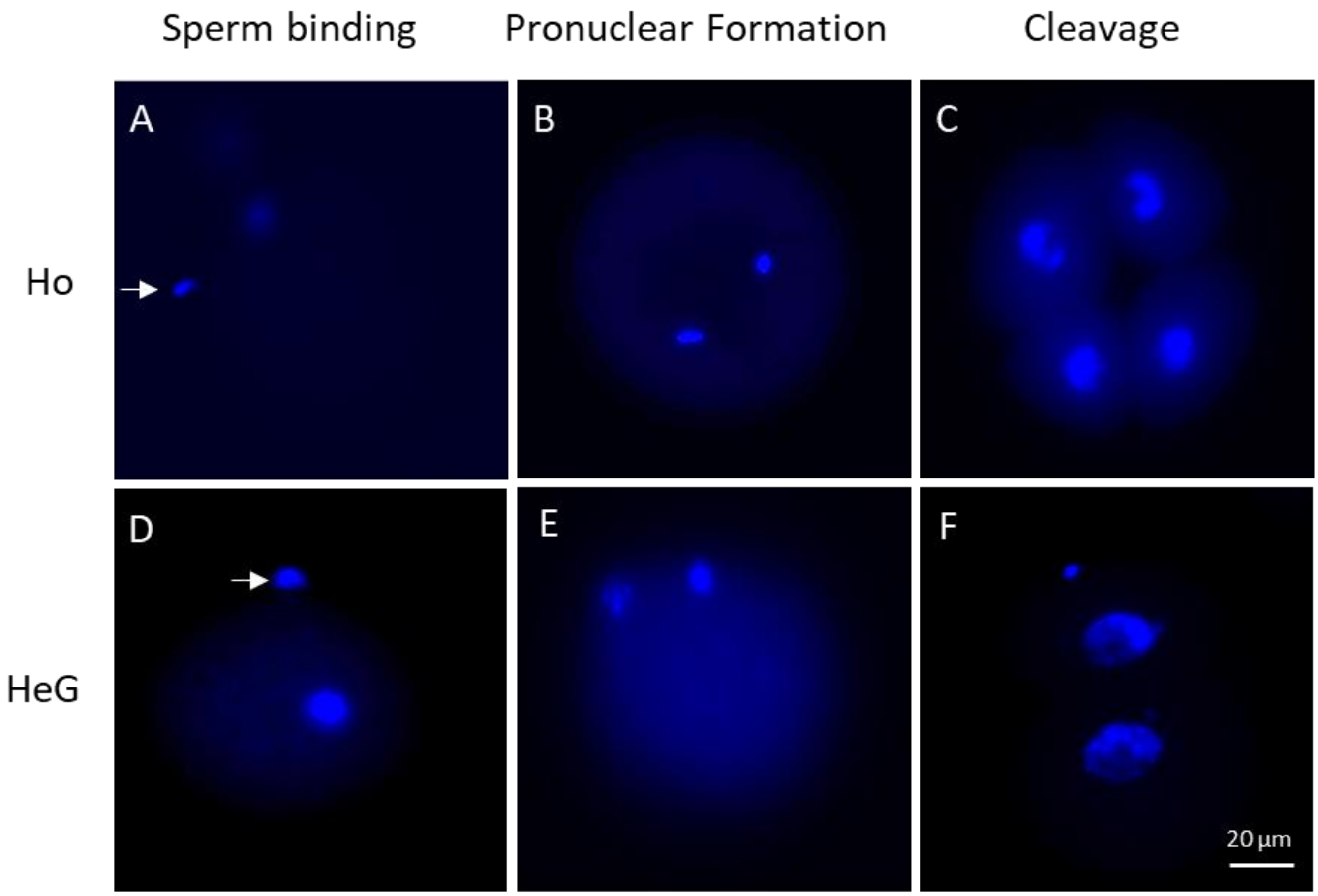


**Fig. S3. Heterologous IVF between bovine oocytes and cat sperm.** Sperm-oocyte binding, pronuclear formation, and cleavage after homologous (Ho) (bovine sperm) and heterologous (He) (cat sperm capacitated in Tyrode's medium (HeG)) IVF. Gametes were stained with Hoechst 33342. Ho: Bound bovine sperm after 2.5 hours of co-incubation with zona-intact bovine oocytes (A); pronuclear formation at 6 hpi (B); and embryo cleavage at 48 hpi (C). HeG: Bound cat sperm after 2.5 hours of co-incubation of the gametes (D); pronuclear formation at 18 hpi (E); and hybrid-embryo cleavage at 48 hpi (F). Arrow points to sperm head chromatin. Images were captured with a 63X objective. Scale bar 20 µm.

methods.

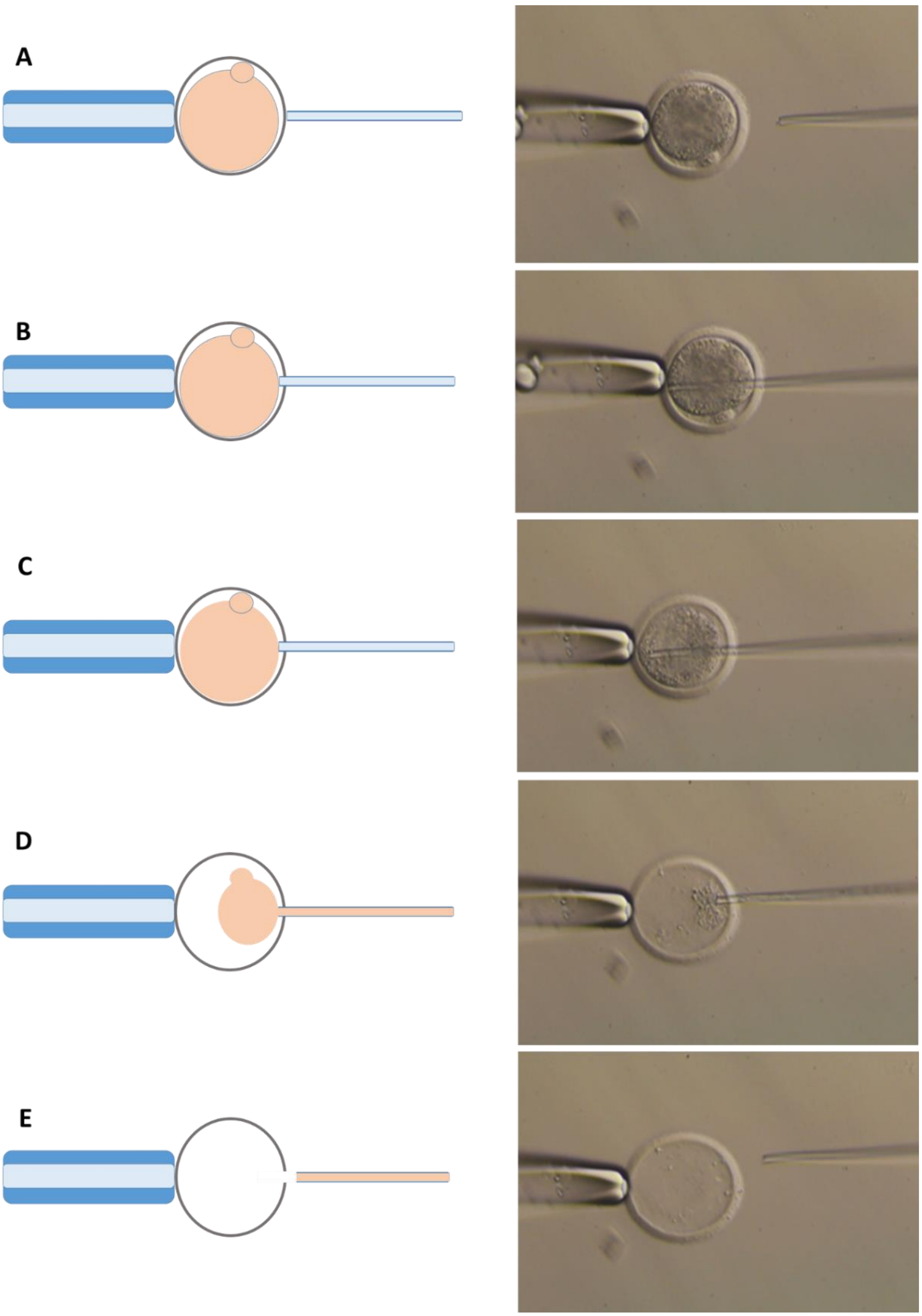


**Fig. S4. Steps (A-E) of the method used to empty the bovine zone pellucida by removing the cytoplasmic contents of the oocyte containing all the organelles, nucleus and membranes, and removing also the polar body.** Details of this procedure can be found in the Materials and methods.

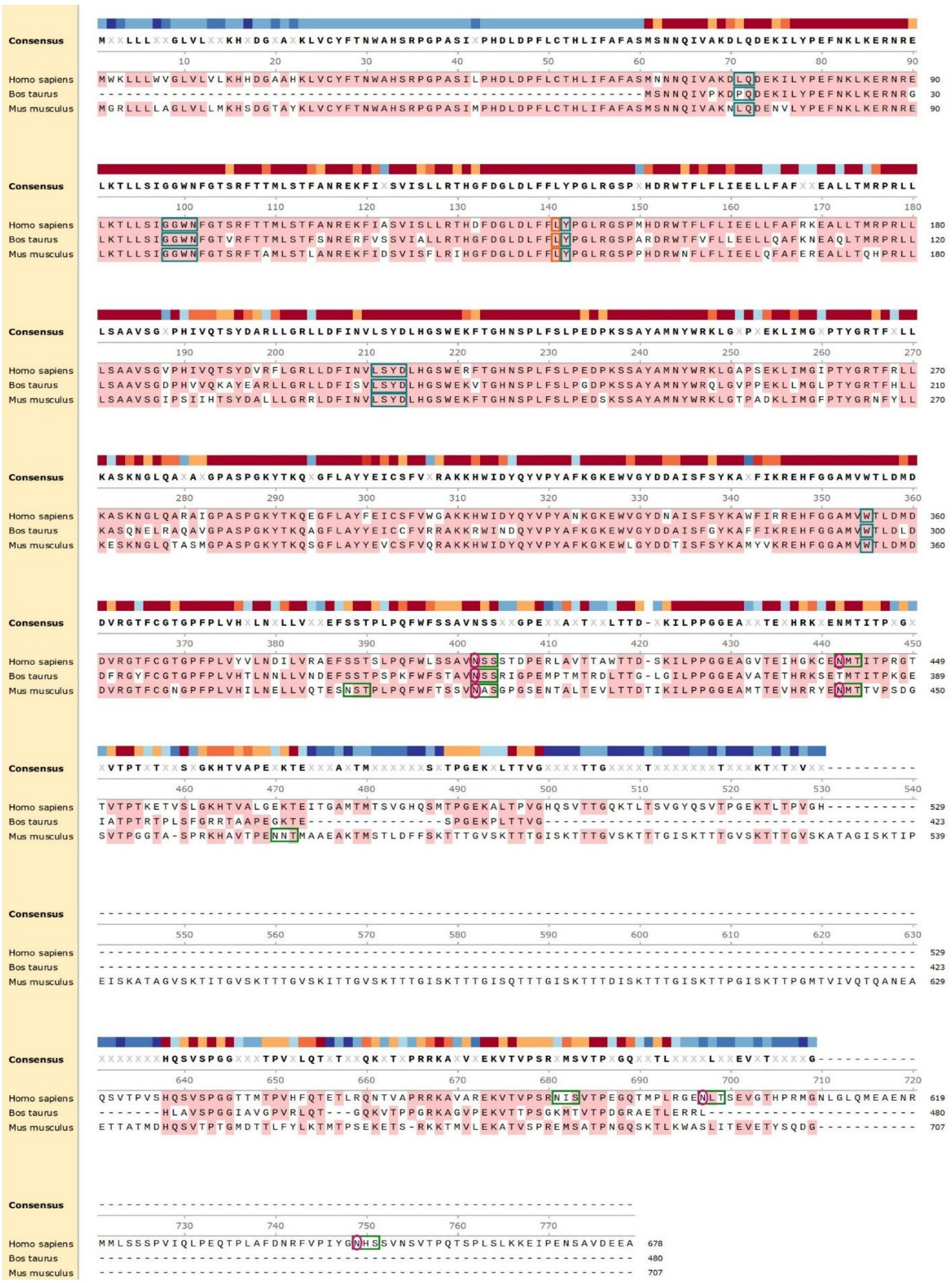

**Fig. S5. Sequence alignments of OVGP1 from Homo sapiens (NCBI AAI36407.1), Bos taurus (NP_001073685.1) and Mus musculus (AAI37996.1)**. Sequences were aligned with the tool

MUSCLE (Multiple Sequence Comparison by Log-Expectation) using Snapgene software. Consensus conserved amino acids are represented in the colored blocks: red = score over the 50% threshold and blue = less conserved. Amino acids corresponding to the consensus are indicated in light pink. N-glycosylation sites were predicted using the NetNGlyc 1.0 Server of DTU Health Tech Bioinformatic Services. The dark green boxes indicate possible asparagine (N) glycosylation. Residues adjacent to N in the sequons Asn-Xaa-Ser/Thr are circled in dark pink. Chitin-binding sites predicted by the UniProt server appear in turquoise boxes. Orange boxes locate the essential glutamic acid of CH18 family members.

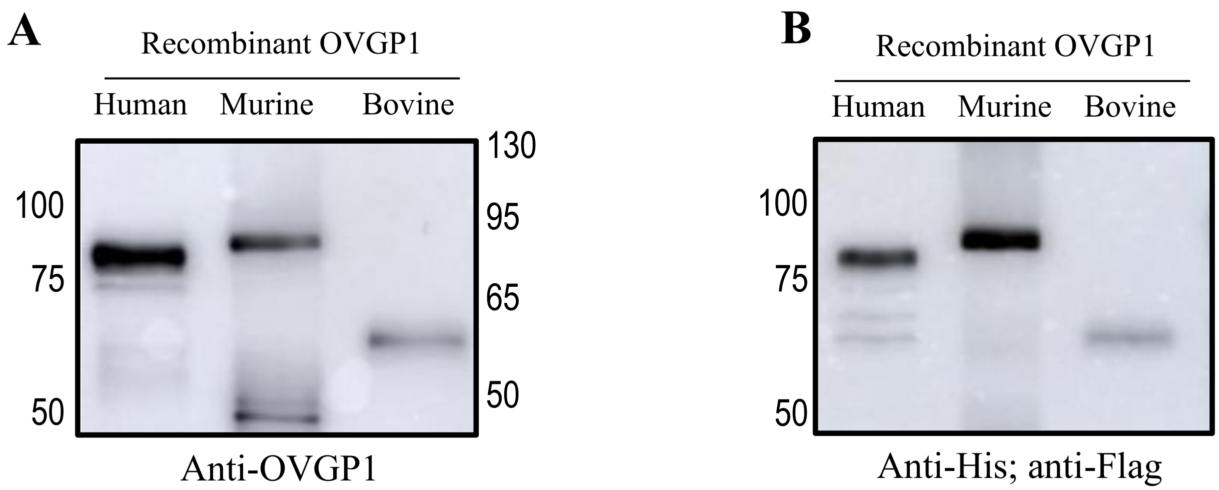


**Fig. S6. Recombinant OVGP1 from human, murine and bovine recognized with anti-OVGP1 (A) and anti-His/anti-Flag (B).** Panel **A** is taken directly from Figure 4 in order to compare antibody recognition. In **B**, is represented the expression of recombinant proteins detected with an anti-His and anti-Flag co-incubated antibodies and developed simultaneously. Flag is detecting hOVGP1 and mOVGP1, while His is detecting bOVGP1.

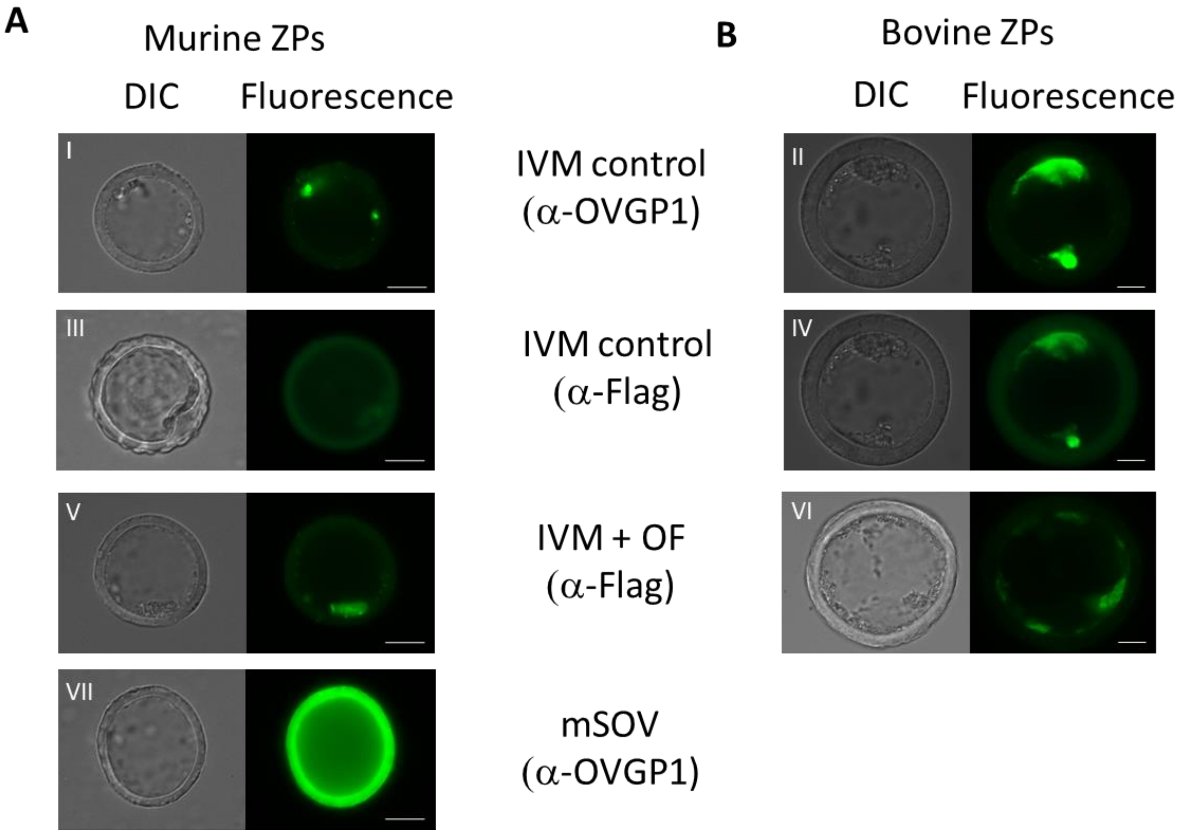


**Fig. S7. Representation of immunofluorescence of control ZPs of IVM murine and bovine oocytes, ZPs incubated with murine or bovine oviductal fluid, and ZPs from murine superovulated (SOV).** IVM murine (**A**) and bovine (**B**) ZPs were fixed and imaged by conventional fluorescence and DIC microscopy using rabbit polyclonal antibody to the human OVGP1 (**I, II**) or a mouse monoclonal antibody against Flag-tag (**III, IV**). IVM ZPs from murine and bovine oocytes were incubated for 30 min with murine or bovine oviductal fluid (respectively) and imaged using anti-Flag tag **(V, VI**). ZPs obtained from oocytes of the oviducts of superovulated mice females (SOV) incubated with anti-OVGP1 (**VII**).

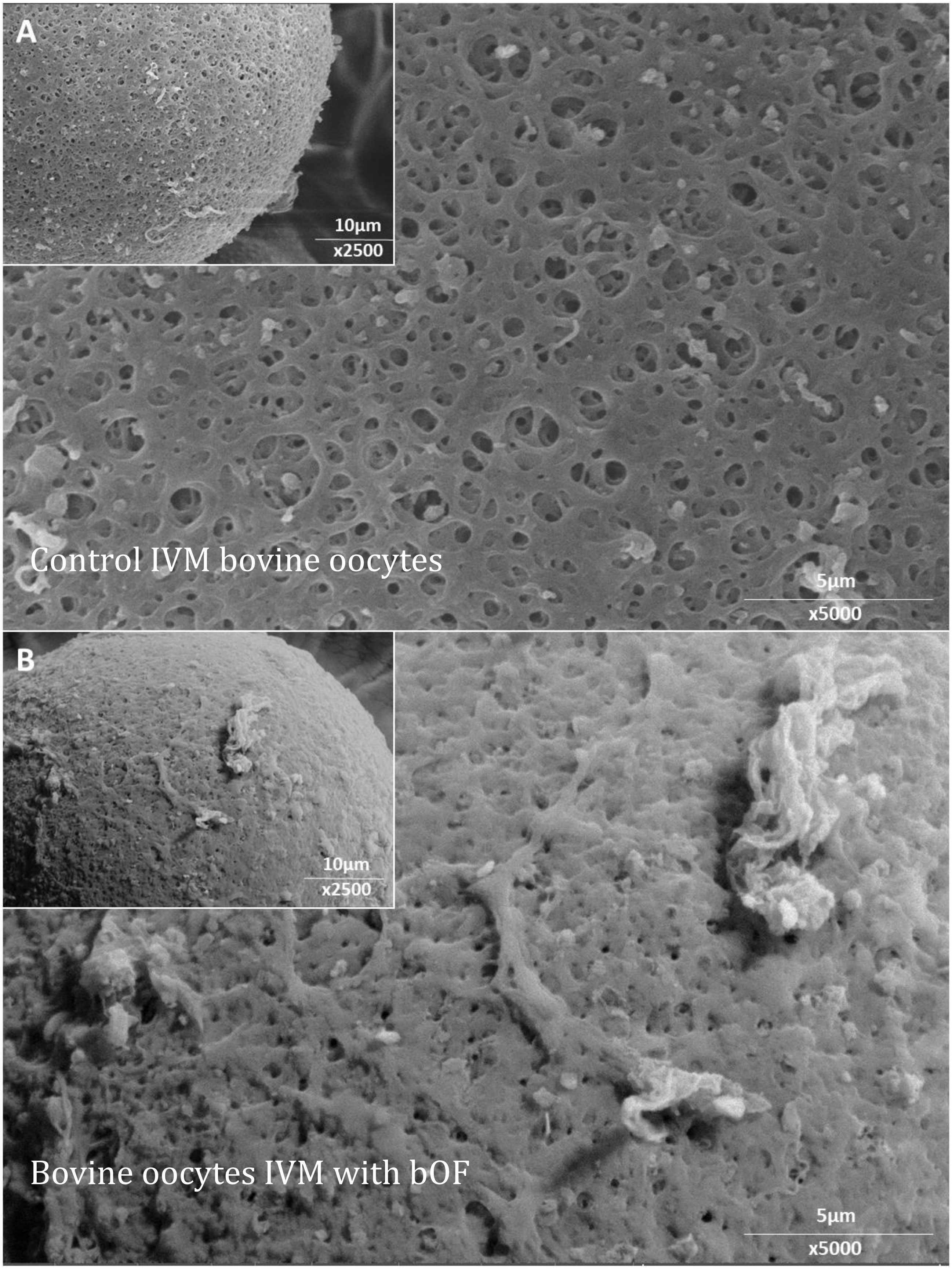
A
10µm
x2500
Control IVM bovine oocytes
5µm
x5000
B
10µm
x2500
Bovine oocytes IVM with bOF
5µm
x5000

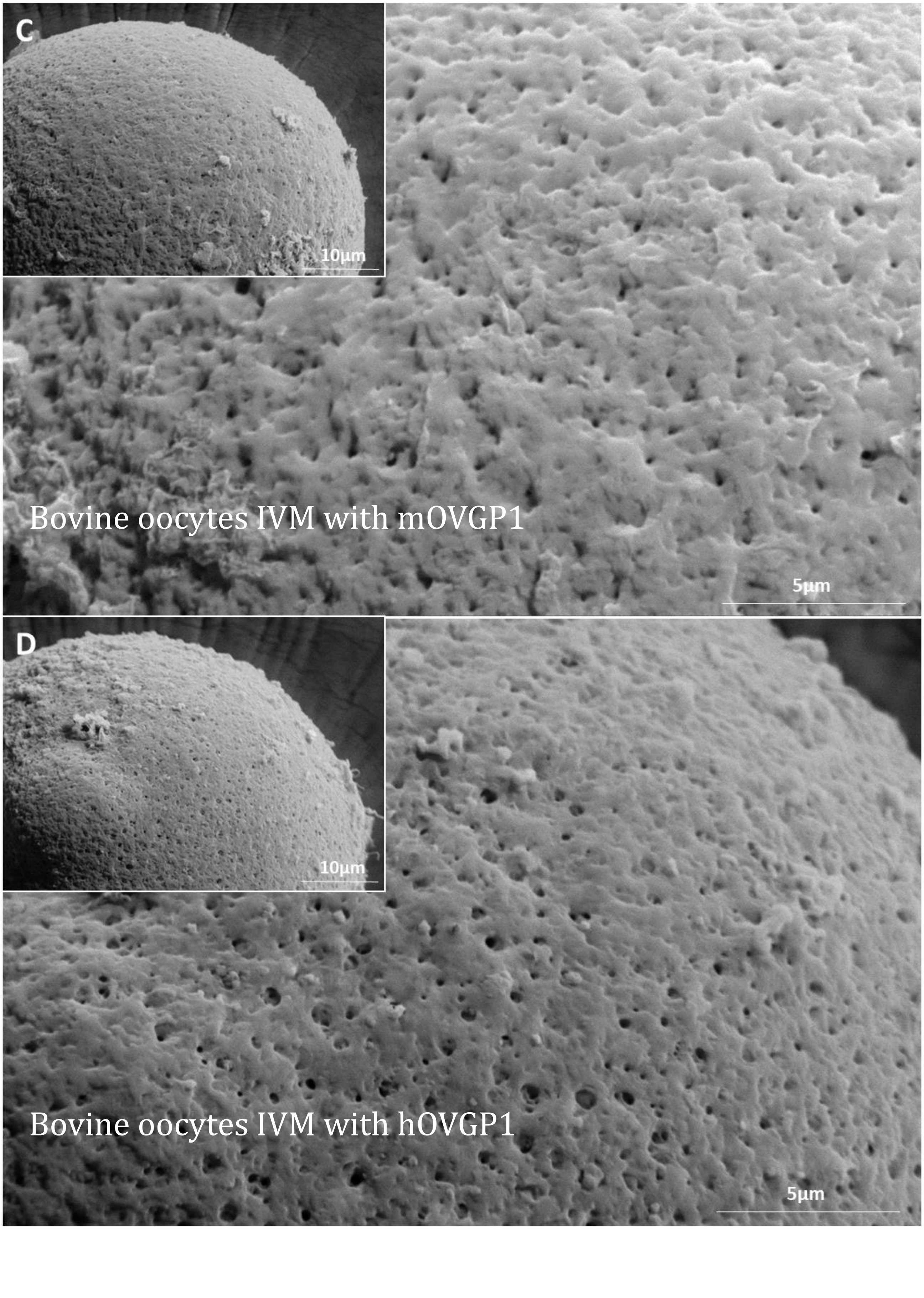
C
10µm
Bovine oocytes IVM with mOVGP1
5µm
D
10µm
Bovine oocytes IVM with hOVGP1
5µm

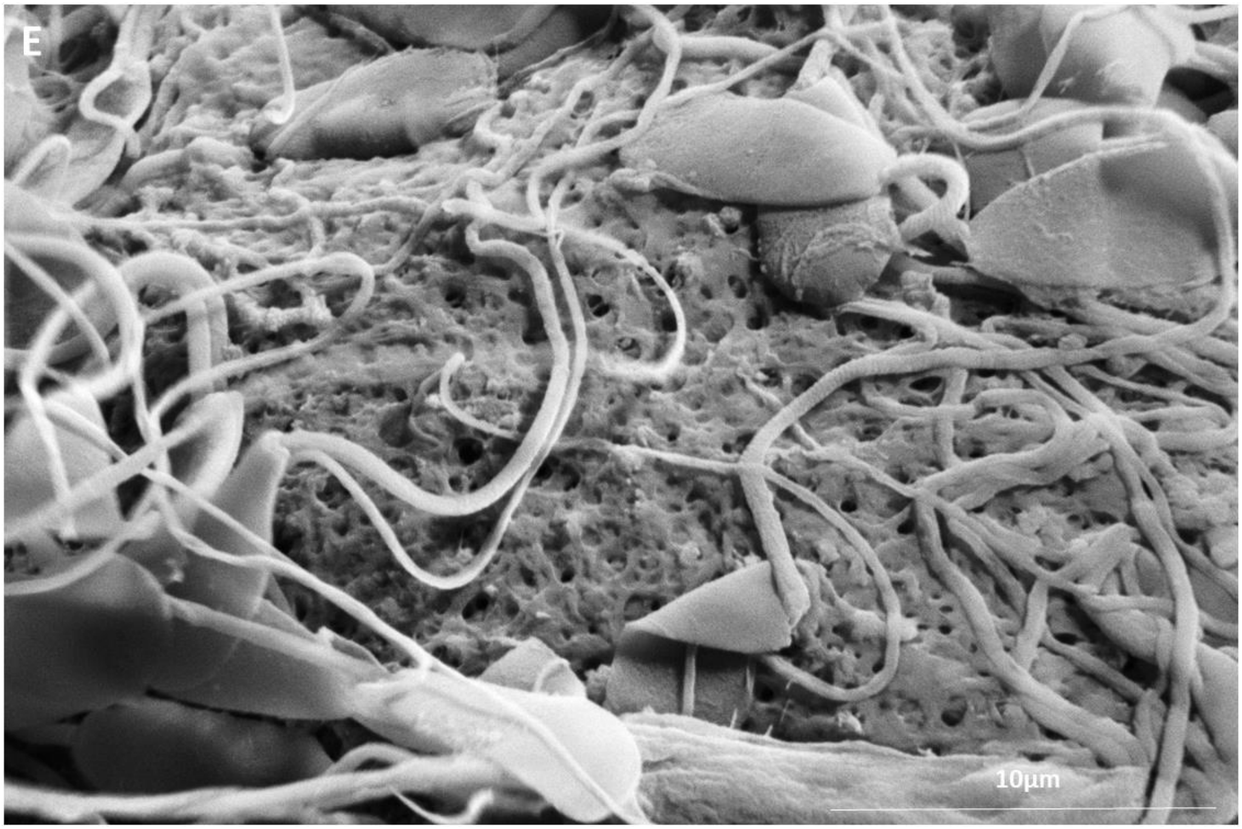


**Fig. S8. Scanning electron micrographs (SEM) of the outer surface of the bovine ZP** treated or not with bovine OF (A, B), mOVGP1 (C), or hOVGP1 (D). (E) Image showing bovine sperm on the surface of the bovine ZP after having been in contact with OVGP1, showing the comparison between the size of the sperm and the size of the pores of the ZP.

**Table S1**. Rates of sperm-ZP binding, pronuclear formation, and cleavage after homologous (bovine sperm) and heterologous (human sperm, media A or B) co-incubation with bovine ovarian oocytes recorded at different times post-insemination

| Semen group (N) | Sperm binding | Pronuclear formation | | | | Cleavage rate |
|---|---|---|---|---|---|---|
| | 2.5 hpi, n | 6 hpi, n (%) | 12 hpi, n (%) | 18 hpi, n (%) | 22 hpi, n (%) | 48 hpi, n (%) |
| Ho (133) | 32 (0.5 ± 0.17) | | | 36 (72.1 ± 7.3)[a] | | 65 (86.0 ± 7.8)[a] |
| HeA (200) | 31 (0.3 ± 0.17) | 32 (34.4 ± 4,7) | 35 (25.5 ± 6.1) | 31 (54.8 ± 7.1)[b] | 30 (56.6 ± 8.1) | 41 (51.2 ± 8.3)[b] |
| HeB (236) | 33 (0.2 ± 0.17) | 36 (30.5 ± 5.4) | 37 (32.1 ± 6.8) | 36 (33.5 ± 7.3)[c] | 38 (34.3 ± 6.6) | 56 (34.4 ± 7.1)[b] |
| Parth (28) | | | | | | 28 (3.3 ± 3.6)[c] |

Ho = homologous IVF with bovine sperm; HeA = heterologous IVF with human sperm using G-IVF™ PLUS medium; HeB = heterologous IVF with human sperm using Fert medium; Parth = parthenogenic non-fertilized oocytes. Sperm binding was expressed as the average number of spermatozoa that remained bound to the ZP. “N” refers to the total number of oocytes fertilized per treatment or non-fertilized oocytes (Parth); “n” refers to the total number of fertilized oocytes/presumptive zygotes or non-fertilized oocytes (Parth) recorded at each time point to determine rates of sperm binding, pronuclear formation, and embryo cleavage. Pronuclear formation was expressed as the percentage of oocytes with two pronuclei at each time point over the number (n) of oocytes examined. Cleavage rate was expressed as the percentage of 2-cell embryos over the number (n) of fertilized or non-fertilized (Parth) oocytes. Analysis was conducted by using a one-way ANOVA, followed by Tukey´s post hoc test. Significance level was set at $p < 0.05$. Different superscripts (a, b, c) in the same column indicate significant differences ($p < 0.05$; n=3); Values are expressed as the mean ± SD.

**Table S2**. Rates of sperm-ZP binding, pronuclear formation, and cleavage after homologous (bovine sperm) and heterologous (murine sperm) co-incubation with bovine ovarian oocytes at different times post-insemination

| Semen group (N) | Sperm binding | Pronuclear formation | Cleavage rate | | |
|---|---|---|---|---|---|
| | 2.5 hpi, n | 18 hpi, n (%) | 20 hpi, n (%) | 26 hpi, n (%) | 48 hpi, n (%) |
| Ho (231) | 46 (0.6 ± 1.0) | 86 (75.8 ± 10.21)[a] | | | 99 (81.8 ± 1.9)[a] |
| He (436) | 47 (0.6 ± 1.0) | 86 (44.3 ± 6.7)[b] | 87 (44.9 ± 8.8) | 85 (44.7 ± 3.6) | 131 (52.6 ± 3.1)[b] |
| Parth (43) | | | | | 43 (4.4 ± 8.7)[c] |

Ho = homologous IVF with bovine sperm; He = heterologous IVF with murine sperm. Parth = parthenogenic non-fertilized oocytes. Sperm binding was expressed as the average number of spermatozoa that remained bound to the ZP. “N” refers to the total number of oocytes fertilized per treatment or non-fertilized oocytes (Parth); “n” refers to the total number of fertilized oocytes/presumptive zygotes or non-fertilized oocytes (Parth) recorded at each time point to determine rates of sperm binding, pronuclear formation, and embryo cleavage. Pronuclear formation was expressed as the percentage of oocytes with two pronuclei at each time point over the number (n) of oocytes examined. Cleavage rate was expressed as the percentage of 2-cell embryos over the number (n) of fertilized or non-fertilized (Parth) oocytes. Analysis was conducted by using a one-way ANOVA, followed by Tukey´s post hoc test. Significance level was set at $p < 0.05$. Different superscripts (a, b, c) in the same column indicate significant differences ($p < 0.05$; n=3); Values are expressed as the mean ± SD.

**Table S3**. Rates of sperm-ZP binding, pronuclear formation, and embryo cleavage after homologous (bovine sperm) and heterologous (cat sperm) co-incubation with bovine ovarian oocytes at different times post-insemination

| Semen group (N) | Sperm binding | Pronuclear formation | | Cleavage |
|---|---|---|---|---|
| | 2.5 hpi, n | 6 hpi, n (%) | 18 hpi, n (%) | 48 hpi, n (%) |
| Ho (184) | 43 (0.5 ± 0.9) | | 62 (71.0 ± 2.3)$^{a}$ | 79 (83.4 ± 1.7)$^{a}$ |
| He (259) | 41 (0.2 ± 0.7) | 48 (31.3 ± 17.14) | 58 (34.1 ± 8.8)$^{b}$ | 112 (41.3 ± 2.1)$^{b}$ |
| Parth (44) | | | | 44 (2.2 ± 6.6)$^{c}$ |

Ho = homologous IVF with bovine sperm; He = heterologous IVF with cat sperm using Tyrode's medium; Parth = parthenogenic non-fertilized oocytes. Sperm binding was expressed as the average number of spermatozoa that remained bound to the ZP. "N" refers to the total number of oocytes fertilized per treatment or non-fertilized oocytes (Parth); "n" refers to the total number of fertilized oocytes/presumptive zygotes or non-fertilized oocytes (Parth) recorded at each time point to determine rates of sperm-ZP binding, pronuclear formation, and cleavage. Pronuclear formation was expressed as the percentage of oocytes with two pronuclei at each time point over the number (n) of oocytes examined. Cleavage rate was expressed as the percentage of 2-cell embryos over the number (n) of fertilized or non-fertilized (Parth) oocytes. Analysis was conducted by using a one-way ANOVA, followed by Tukey´s post hoc test. Significance level was set at $p < 0.05$. Different superscripts (a, b, c) in the same column indicate significant differences ($p < 0.05$; n=3); Values are expressed as the mean ± SD.

**Table S4.** Rates of sperm-ZP binding, pronuclear formation, and cleavage after homologous (bovine sperm) and heterologous (human sperm, media A or B) co-incubation with bovine ovarian oocytes (previously incubated with oviductal fluid for 30 minutes) at different times post-insemination

| Semen group | Sperm binding | Pronuclear formation | | | Cleavage | |
|---|---|---|---|---|---|---|
| (N) | 2.5 hpi n | 6 hpi n (%) | 12 hpi n (%) | 18 hpi n (%) | 22 hpi n (%) | 48 hpi n (%) |
| Ho (135) | 32 (0.5 ± 0.9)[a] | | | 33 (66.7 ± 3.6)[a] | | 70 (85.7 ± 3.4)[a] |
| HeA (267) | 34 (0.01 ± 0.2)[b] | 38 (0.0 ± 0.0 | 43 (0.0 ± 0.0) | 49 (0.0 ± 0.0)[b] | 44 (0.0 ± 0.0) | 59 (3.3 ± 2.6)[b] |
| HeB (235) | 31 (0.03 ± 0.2)[b] | 36 (0.0 ± 0.0 | 34 (0.0 ± 0.0) | 36 (0.0 ± 0.0)[b] | 36 (0.0 ± 0.0) | 62 (3.1 ± 4.5)[b] |
| Parth (46) | | | | | | 28 (1.8 ± 4.1)[b] |

Ho = homologous IVF with bovine sperm; HeA = heterologous IVF with human sperm in IVF-G medium; HeB = heterologous IVF with human sperm in Fert medium; Parth = parthenogenic non-fertilized oocytes. "N" refers to the total number of oocytes fertilized per treatment or non-fertilized oocytes (Parth); "n" refers to the total number of fertilized oocytes/presumptive zygotes or non-fertilized oocytes (Parth) recorded at each time point to determine rates of sperm-ZP binding, pronuclear formation and cleavage. Sperm binding was expressed as the average number of spermatozoa that remained bound to the ZP. Pronuclear formation was expressed as the percentage of oocytes with two pronuclei at each time point over the number (n) of oocytes examined. Cleavage rate was expressed as the percentage of 2-cell embryos over the number (n) of fertilized or non fertilized (Parth) oocytes. Analysis was conducted by using a one-way ANOVA, followed by Tukey´s post hoc test. Different superscripts (a, b) in the same column indicate significant differences ($p < 0.05$; n=3). Values are expressed as the mean ± SD.

**Table S5**. Rates of sperm penetration and binding to the murine ZP after the homologous (murine sperm) and heterologous (bovine sperm) EZPT using ZPs prepared from oocytes from murine ovaries (HoA, HeA) and from the oviducts of superovulated female mice (HoB, HeB)

| Group<br>N | Sperm | ZPs from murine oocytes | TMC | Penetrated ZPs (%) | Sperm penetration inside ZP (mean) | Max. no. of sperm inside a single ZP | ZP with bound sperm (%) | Sperm bound to ZPs (mean) | Max. no. of sperm bound to a single ZP |
|---|---|---|---|---|---|---|---|---|---|
| HoA<br>56 | Murine | Ovary | 8.76 ± 1.07 | 56<br>(100 ± 0)[a] | 848<br>(15.08 ± 1.19)[a] | 29 ± 5.50[a] | 56<br>(100 ± 0)[a] | 1042<br>(19.02 ± 2.21)[a] | 32 ± 2.06[a] |
| HeAb<br>97 | Bovine | Ovary | 3.67 ± 0.19 | 56<br>(57.85 ± 5.45)[b] | 167<br>(2.98 ± 0.02)[b] | 10 ± 1.4[b] | 91<br>(93.87 ± 2.65)[b] | 303<br>(3.33 ± 0.52)[b] | 9 ±1.42[b] |
| HeAh<br>85 | Human | Ovary | 6.12 ± 0.17 | 60<br>(70.83 ± 5.89)[b] | 177<br>(2.95 ± 0,07)[b] | 9 ± 0.71[b] | 79<br>(92.92 ± 0.59)[b] | 263<br>(3.34 ± 0.35)[b] | 11 ± 1.41[b] |
| HoB<br>37 | Murine | Oviduct | 8.76 ± 1.07 | 37<br>(100 ± 0)[a] | 423<br>(23.86 ± 0.79)[a] | 18 ± 4.24[a] | 37<br>(100 ± 0)[a] | 645<br>(10.02 ± 1.81)[a] | 28 ± 2.83[a] |
| HeBb<br>131 | Bovine | Oviduct | 3.67 ± 0.19 | 3<br>(6.08± 1.36)[c] | 3<br>(1 ± 0)[c] | 1 ± 0[c] | 21<br>(21.46 ± 2.06)[c] | 62<br>(2.96 ± 0.33)[c] | 4 ± 0.71[c] |
| HeBh<br>65 | Human | Oviduct | 6.12 ± 0.17 | 3<br>(4.72 ± 0,39)[c] | 3<br>(0.05 ± 0.01)[c] | 1 ± 0[c] | 14<br>(22.50 ± 3.54)[c] | 39<br>(2.70 ± 0.42)[c] | 5 ± 2.83[c] |

HoA = homologous EZPT with murine sperm (oocytes obtained from ovary). HeAb = heterologous EZPT with bovine sperm (oocytes obtained from ovary). HeAh = heterologous EZPT with human sperm (oocytes obtained from ovary). HoB = homologous EZPT with murine sperm (oocytes obtained from the oviduct of SOV females). HeBb = heterologous EZPT with bovine sperm (oocytes obtained from the oviduct of SOV females). HeBh = heterologous EZPT with human sperm (oocytes obtained from the oviduct of SOV females). TMC = total motile sperm count after capacitation calculated by multiplying the volume by concentration (million sperm/mL) and by the motility (%) after capacitation. Penetrated ZPs refers to the mean number of ZPs penetrated by at least one sperm. ZPs with bound sperm was calculated as the mean number of ZP with, at least, one bound sperm. Sperm penetration was taken as the total number of sperm observed inside the zonas. Bound sperm was expressed as the mean number of sperm that remained bound to the ZP. “N” refers to the total number of ZPs fertilized per treatment. Analysis was conducted by using a one-way ANOVA, followed by Tukey´s post hoc test. Different superscripts (a, b, c) in the same column indicate significant differences ($p < 0.05$; n=3). Values are expressed as the mean ± SD.

**Table S6.** Rates of sperm penetration and binding to the bovine ZP after homologous (bovine sperm) and heterologous (human or murine sperm) EZPT using bovine ZPs co-incubated without bovine oviductal fluid (HoA, HeA) or with bovine oviductal fluid (HoB, HeB)

| Group N | Sperm | Oviductal fluid | TMC | Penetrated ZPs (%) | Sperm penetration inside ZP (mean) | Max. no. of sperm inside a single ZP | ZP with bound sperm (%) | Sperm bound to ZPs (mean) | Max. no. of sperm bound to a single ZP |
|---|---|---|---|---|---|---|---|---|---|
| HoA 90 | Bovine | - | 4.16 ± 0.27 | 90 (100 ± 0)[a] | 853 (9.13 ± 1.23)[a] | 30 ± 1.41[a] | 90 (100 ± 0)[a] | 830 (9.27± 0.58)[a] | 23 ± 1.12[a] |
| HeAh 406 | Human | - | 9.14 ± 3.98 | 283 (69.7± 12.1)[b] | 1588 (5.61 ± 2.83)[b] | 23 ± 4.2[b] | 96 (23.65 ± 8.59)[b] | 188 (1.49 ± 0.24)[b] | 6 ± 1.27[b] |
| HeAm 83 | Murine | - | 8.76 ± 1.07 | 80 (96.25 ± 5.3)[a] | 300 (3.73 ± 0.31)[C] | 21 ± 2.12[b] | 83 (100 ± 0)[a] | 490 (5.92 ± 0.81)[c] | 20 ± 3.54[a] |
| HoB 56 | Bovine | + | 4.16 ± 0.27 | 56 (100 ± 0)[a] | 585 (10.52 ± 0.56)[a] | 24 ± 2.82[b] | 56 (100 ± 0)[a] | 574 (10.02 ± 1.81)[a] | 23 ± 2.12[a] |
| HeBh 131 | Human | + | 9.14 ± 3.98 | 0 (0 ± 0)[c] | 0 (0± 0)[c] | 0 ± 0[c] | 10 (7.42 ± 2.53)[c] | 10 (1 ± 0)[d] | 1 ± 0[c] |
| HeBm 48 | Murine | + | 9.71 ± 0.61 | 4 (0.08 ± 0.005)[c] | 4 (1 ± 0)[c] | 1 ± 0[c] | 20 (41.74 ± 2.46)[b] | 45 (0.94 ± 0.08)[d] | 5 ± 0.71[b] |

HoA = homologous EZPT with bovine sperm. HeAh = heterologous EZPT with human sperm. HeAm = heterologous EZPT with murine sperm. HoB = homologous EZPT with bovine sperm (ZPs were co-incubated with bovine oviductal fluid for 30 minutes). HeBh = heterologous EZPT with human sperm (ZPs were co-incubated with bovine oviductal fluid for 30 minutes). HeBm = heterologous EZPT with murine sperm (ZPs were co-incubated with bovine oviductal fluid for 30 minutes). TMC = total motile sperm count after capacitation calculated by multiplying the volume by concentration (million sperm/mL) and by the motility (%) after capacitation. Penetrated ZPs refers to the mean number of ZPs penetrated by at least one sperm. ZPs with bound sperm was calculated as the mean number of ZPs with, at least, one bound sperm. Sperm penetration was taken as the total number of sperm observed inside the zonas. Bound sperm was expressed as the mean number of sperm that remained bound to the ZP. “N” refers to the total number of ZPs fertilized per treatment. Analysis was conducted by using a one-way ANOVA, followed by Tukey´s post hoc test. Different superscripts (a, b, c, d) in the same column indicate significant differences ($p < 0.05$; n=3). Values are expressed as the mean ± SD.

**Table S7.** O-glycosylation prediction sites in human, mouse and bull OVGP1. Identification and location of the GalNAc-type O-glycosylation sites with the online server NetOGlyc 4.0 (*1*)

| | Start | End | Score | Comment | | Start | End | Score | Comment | | Start | End | Score | Comment |
|---|---|---|---|---|---|---|---|---|---|---|---|---|---|---|
| HOMO | 29 | 29 | 0.148743 | | MUS | 17 | 17 | 0.0664484 | | BOS | 2 | 2 | 0.477639 | |
| HOMO | 34 | 34 | 0.513945 | + | MUS | 20 | 20 | 0.0136154 | | BOS | 33 | 33 | 0.0141907 | |
| HOMO | 40 | 40 | 0.133798 | | MUS | 29 | 29 | 0.13254 | | BOS | 36 | 36 | 0.0521786 | |
| HOMO | 52 | 52 | 0.0144943 | | MUS | 34 | 34 | 0.5 | + | BOS | 44 | 44 | 0.211234 | |
| HOMO | 60 | 60 | 0.0428766 | | MUS | 40 | 40 | 0.153781 | | BOS | 48 | 48 | 0.201186 | |
| HOMO | 93 | 93 | 0.0158923 | | MUS | 52 | 52 | 0.0118199 | | BOS | 49 | 49 | 0.283351 | |
| HOMO | 96 | 96 | 0.0261905 | | MUS | 60 | 60 | 0.0436814 | | BOS | 52 | 52 | 0.124074 | |
| HOMO | 104 | 104 | 0.216291 | | MUS | 62 | 62 | 0.0894996 | | BOS | 53 | 53 | 0.100269 | |
| HOMO | 105 | 105 | 0.252093 | | MUS | 93 | 93 | 0.0153551 | | BOS | 55 | 55 | 0.207505 | |
| HOMO | 108 | 108 | 0.14834 | | MUS | 96 | 96 | 0.0240849 | | BOS | 62 | 62 | 0.226985 | |
| HOMO | 109 | 109 | 0.315057 | | MUS | 104 | 104 | 0.143094 | | BOS | 63 | 63 | 0.0478178 | |
| HOMO | 112 | 112 | 0.180764 | | MUS | 105 | 105 | 0.199945 | | BOS | 70 | 70 | 0.0242045 | |
| HOMO | 113 | 113 | 0.114855 | | MUS | 108 | 108 | 0.111836 | | BOS | 88 | 88 | 0.081231 | |
| HOMO | 123 | 123 | 0.030659 | | MUS | 112 | 112 | 0.0957522 | | BOS | 95 | 95 | 0.0428404 | |
| HOMO | 126 | 126 | 0.0314989 | | MUS | 113 | 113 | 0.100753 | | BOS | 114 | 114 | 0.078206 | |
| HOMO | 130 | 130 | 0.0220408 | | MUS | 123 | 123 | 0.00798697 | | BOS | 122 | 122 | 0.240522 | |
| HOMO | 148 | 148 | 0.0445812 | | MUS | 126 | 126 | 0.0128272 | | BOS | 126 | 126 | 0.474913 | |
| HOMO | 155 | 155 | 0.0372563 | | MUS | 148 | 148 | 0.0325529 | | BOS | 149 | 149 | 0.0380635 | |
| HOMO | 174 | 174 | 0.0488918 | | MUS | 174 | 174 | 0.0484669 | | BOS | 152 | 152 | 0.00857951 | |
| HOMO | 182 | 182 | 0.160501 | | MUS | 182 | 182 | 0.166503 | | BOS | 158 | 158 | 0.0631895 | |
| HOMO | 186 | 186 | 0.462315 | | MUS | 186 | 186 | 0.388758 | | BOS | 163 | 163 | 0.275655 | |
| HOMO | 194 | 194 | 0.11187 | | MUS | 190 | 190 | 0.0693216 | | BOS | 167 | 167 | 0.487753 | |
| HOMO | 195 | 195 | 0.0571954 | | MUS | 194 | 194 | 0.0973003 | | BOS | 171 | 171 | 0.482996 | |
| HOMO | 212 | 212 | 0.00914751 | | MUS | 195 | 195 | 0.057626 | | BOS | 178 | 178 | 0.0814828 | |
| HOMO | 218 | 218 | 0.0565867 | | MUS | 212 | 212 | 0.00829588 | | BOS | 179 | 179 | 0.348292 | |
| HOMO | 223 | 223 | 0.195033 | | MUS | 218 | 218 | 0.047986 | | BOS | 202 | 202 | 0.139565 | |
| HOMO | 227 | 227 | 0.375705 | | MUS | 223 | 223 | 0.18869 | | BOS | 206 | 206 | 0.199712 | |
| HOMO | 231 | 231 | 0.310035 | | MUS | 227 | 227 | 0.246007 | | BOS | 213 | 213 | 0.5 | + |
| HOMO | 238 | 238 | 0.115751 | | MUS | 231 | 231 | 0.246397 | | BOS | 226 | 226 | 0.696097 | + |
| HOMO | 239 | 239 | 0.335736 | | MUS | 236 | 236 | 0.17176 | | BOS | 231 | 231 | 0.428867 | |
| HOMO | 253 | 253 | 0.0093353 | | MUS | 238 | 238 | 0.102913 | | BOS | 276 | 276 | 0.0105995 | |
| HOMO | 262 | 262 | 0.09816 | | MUS | 239 | 239 | 0.289418 | | BOS | 296 | 296 | 0.0180258 | |
| HOMO | 266 | 266 | 0.142253 | | MUS | 251 | 251 | 0.029901 | | BOS | 309 | 309 | 0.0717533 | |
| HOMO | 273 | 273 | 0.4854 | | MUS | 262 | 262 | 0.133467 | | BOS | 317 | 317 | 0.183042 | |
| HOMO | 286 | 286 | 0.829831 | + | MUS | 273 | 273 | 0.193716 | | BOS | 328 | 328 | 0.222882 | |
| HOMO | 291 | 291 | 0.174597 | | MUS | 279 | 279 | 0.662371 | + | BOS | 329 | 329 | 0.779465 | + |

| | | | | |
|---|---|---|---|---|
| HOMO | 304 | 304 | 0.0267057 | |
| HOMO | 336 | 336 | 0.0314593 | |
| HOMO | 338 | 338 | 0.0119623 | |
| HOMO | 356 | 356 | 0.0239896 | |
| HOMO | 365 | 365 | 0.131311 | |
| HOMO | 369 | 369 | 0.120416 | |
| HOMO | 388 | 388 | 0.491023 | |
| HOMO | 389 | 389 | 0.824032 | + |
| HOMO | 390 | 390 | 0.6421 | + |
| HOMO | 391 | 391 | 0.739353 | + |
| HOMO | 398 | 398 | 0.757269 | + |
| HOMO | 399 | 399 | 0.872818 | + |
| HOMO | 403 | 403 | 0.797847 | + |
| HOMO | 404 | 404 | 0.793866 | + |
| HOMO | 405 | 405 | 0.955482 | + |
| HOMO | 406 | 406 | 0.740014 | + |
| HOMO | 414 | 414 | 0.978317 | + |
| HOMO | 415 | 415 | 0.912469 | + |
| HOMO | 418 | 418 | 0.926281 | + |
| HOMO | 419 | 419 | 0.913712 | + |
| HOMO | 421 | 421 | 0.780885 | + |
| HOMO | 433 | 433 | 0.403671 | |
| HOMO | 443 | 443 | 0.803539 | + |
| HOMO | 445 | 445 | 0.760132 | + |
| HOMO | 449 | 449 | 0.950001 | + |
| HOMO | 450 | 450 | 0.883589 | + |
| HOMO | 452 | 452 | 0.897374 | + |
| HOMO | 454 | 454 | 0.952252 | + |
| HOMO | 457 | 457 | 0.844377 | + |
| HOMO | 459 | 459 | 0.950975 | + |
| HOMO | 464 | 464 | 0.852675 | + |
| HOMO | 471 | 471 | 0.696304 | + |
| HOMO | 474 | 474 | 0.846582 | + |
| HOMO | 478 | 478 | 0.776336 | + |
| HOMO | 480 | 480 | 0.745116 | + |
| HOMO | 481 | 481 | 0.794363 | + |
| HOMO | 486 | 486 | 0.819105 | + |
| HOMO | 488 | 488 | 0.884396 | + |
| HOMO | 495 | 495 | 0.723935 | + |
| HOMO | 501 | 501 | 0.820028 | + |
| HOMO | 503 | 503 | 0.97727 | + |
| HOMO | 504 | 504 | 0.886534 | + |

| | | | | |
|---|---|---|---|---|
| MUS | 281 | 281 | 0.959249 | + |
| MUS | 286 | 286 | 0.791757 | + |
| MUS | 291 | 291 | 0.32144 | |
| MUS | 294 | 294 | 0.286168 | |
| MUS | 304 | 304 | 0.048395 | |
| MUS | 334 | 334 | 0.00966718 | |
| MUS | 336 | 336 | 0.0104758 | |
| MUS | 338 | 338 | 0.0196048 | |
| MUS | 356 | 356 | 0.0175912 | |
| MUS | 365 | 365 | 0.035171 | |
| MUS | 385 | 385 | 0.265295 | |
| MUS | 387 | 387 | 0.549473 | + |
| MUS | 389 | 389 | 0.61239 | + |
| MUS | 390 | 390 | 0.360483 | |
| MUS | 398 | 398 | 0.76942 | + |
| MUS | 399 | 399 | 0.793433 | + |
| MUS | 400 | 400 | 0.887109 | + |
| MUS | 404 | 404 | 0.813503 | + |
| MUS | 408 | 408 | 0.793101 | + |
| MUS | 411 | 411 | 0.761464 | + |
| MUS | 414 | 414 | 0.725498 | + |
| MUS | 418 | 418 | 0.690982 | + |
| MUS | 419 | 419 | 0.677789 | + |
| MUS | 421 | 421 | 0.746303 | + |
| MUS | 433 | 433 | 0.757383 | + |
| MUS | 434 | 434 | 0.316116 | |
| MUS | 444 | 444 | 0.751447 | + |
| MUS | 445 | 445 | 0.394782 | |
| MUS | 448 | 448 | 0.787216 | + |
| MUS | 451 | 451 | 0.5 | + |
| MUS | 453 | 453 | 0.541804 | + |
| MUS | 457 | 457 | 0.835698 | + |
| MUS | 459 | 459 | 0.749447 | + |
| MUS | 466 | 466 | 0.72813 | + |
| MUS | 471 | 471 | 0.756103 | + |
| MUS | 478 | 478 | 0.539962 | + |
| MUS | 480 | 480 | 0.748887 | + |
| MUS | 481 | 481 | 0.616459 | + |
| MUS | 486 | 486 | 0.774382 | + |
| MUS | 488 | 488 | 0.664138 | + |
| MUS | 489 | 489 | 0.858047 | + |
| MUS | 490 | 490 | 0.843136 | + |

| | | | | |
|---|---|---|---|---|
| BOS | 330 | 330 | 0.813643 | + |
| BOS | 332 | 332 | 0.728598 | + |
| BOS | 338 | 338 | 0.930032 | + |
| BOS | 339 | 339 | 0.896458 | + |
| BOS | 343 | 343 | 0.821513 | + |
| BOS | 344 | 344 | 0.892312 | + |
| BOS | 352 | 352 | 0.862842 | + |
| BOS | 354 | 354 | 0.816318 | + |
| BOS | 358 | 358 | 0.767588 | + |
| BOS | 359 | 359 | 0.60832 | + |
| BOS | 373 | 373 | 0.426953 | |
| BOS | 375 | 375 | 0.726137 | + |
| BOS | 379 | 379 | 0.817664 | + |
| BOS | 381 | 381 | 0.78998 | + |
| BOS | 383 | 383 | 0.829896 | + |
| BOS | 385 | 385 | 0.746896 | + |
| BOS | 392 | 392 | 0.65328 | + |
| BOS | 394 | 394 | 0.763928 | + |
| BOS | 396 | 396 | 0.718114 | + |
| BOS | 399 | 399 | 0.571659 | + |
| BOS | 404 | 404 | 0.875147 | + |
| BOS | 411 | 411 | 0.852652 | + |
| BOS | 413 | 413 | 0.827119 | + |
| BOS | 420 | 420 | 0.640815 | + |
| BOS | 421 | 421 | 0.566064 | + |
| BOS | 428 | 428 | 0.77777 | + |
| BOS | 441 | 441 | 0.403374 | |
| BOS | 446 | 446 | 0.770962 | + |
| BOS | 459 | 459 | 0.760108 | + |
| BOS | 460 | 460 | 0.831493 | + |
| BOS | 462 | 462 | 0.757798 | + |
| BOS | 466 | 466 | 0.653766 | + |
| BOS | 468 | 468 | 0.823171 | + |
| BOS | 475 | 475 | 0.116021 | |

| | | | | |
|---|---|---|---|---|
| HOMO | 508 | 508 | 0.780642 | + |
| HOMO | 510 | 510 | 0.739161 | + |
| HOMO | 511 | 511 | 0.895653 | + |
| HOMO | 516 | 516 | 0.863073 | + |
| HOMO | 518 | 518 | 0.913466 | + |
| HOMO | 523 | 523 | 0.767302 | + |
| HOMO | 525 | 525 | 0.753387 | + |
| HOMO | 531 | 531 | 0.808482 | + |
| HOMO | 533 | 533 | 0.898742 | + |
| HOMO | 536 | 536 | 0.828278 | + |
| HOMO | 539 | 539 | 0.891049 | + |
| HOMO | 541 | 541 | 0.95368 | + |
| HOMO | 545 | 545 | 0.897366 | + |
| HOMO | 546 | 546 | 0.894875 | + |
| HOMO | 548 | 548 | 0.726574 | + |
| HOMO | 554 | 554 | 0.7768 | + |
| HOMO | 556 | 556 | 0.938298 | + |
| HOMO | 561 | 561 | 0.917442 | + |
| HOMO | 575 | 575 | 0.93519 | + |
| HOMO | 578 | 578 | 0.910072 | + |
| HOMO | 582 | 582 | 0.93418 | + |
| HOMO | 584 | 584 | 0.85508 | + |
| HOMO | 589 | 589 | 0.855402 | + |
| HOMO | 598 | 598 | 0.640589 | + |
| HOMO | 599 | 599 | 0.929108 | + |
| HOMO | 603 | 603 | 0.806488 | + |
| HOMO | 623 | 623 | 0.774533 | + |
| HOMO | 624 | 624 | 0.894222 | + |
| HOMO | 625 | 625 | 0.890486 | + |
| HOMO | 634 | 634 | 0.826692 | + |
| HOMO | 650 | 650 | 0.733632 | + |
| HOMO | 651 | 651 | 0.915455 | + |
| HOMO | 654 | 654 | 0.858169 | + |
| HOMO | 656 | 656 | 0.926624 | + |
| HOMO | 659 | 659 | 0.87521 | + |
| HOMO | 660 | 660 | 0.942906 | + |
| HOMO | 663 | 663 | 0.864385 | + |
| HOMO | 672 | 672 | 0.582959 | + |

| | | | | |
|---|---|---|---|---|
| MUS | 493 | 493 | 0.871285 | + |
| MUS | 495 | 495 | 0.77954 | + |
| MUS | 496 | 496 | 0.965151 | + |
| MUS | 497 | 497 | 0.917272 | + |
| MUS | 500 | 500 | 0.91103 | + |
| MUS | 502 | 502 | 0.736343 | + |
| MUS | 503 | 503 | 0.961134 | + |
| MUS | 504 | 504 | 0.901609 | + |
| MUS | 507 | 507 | 0.921979 | + |
| MUS | 509 | 509 | 0.729352 | + |
| MUS | 510 | 510 | 0.950811 | + |
| MUS | 511 | 511 | 0.896131 | + |
| MUS | 514 | 514 | 0.889054 | + |
| MUS | 516 | 516 | 0.727671 | + |
| MUS | 517 | 517 | 0.975857 | + |
| MUS | 518 | 518 | 0.936021 | + |
| MUS | 521 | 521 | 0.906595 | + |
| MUS | 523 | 523 | 0.623161 | + |
| MUS | 524 | 524 | 0.94394 | + |
| MUS | 525 | 525 | 0.898366 | + |
| MUS | 528 | 528 | 0.923861 | + |
| MUS | 531 | 531 | 0.871652 | + |
| MUS | 535 | 535 | 0.672261 | + |
| MUS | 537 | 537 | 0.382735 | |
| MUS | 542 | 542 | 0.777675 | + |
| MUS | 545 | 545 | 0.908425 | + |
| MUS | 549 | 549 | 0.882743 | + |
| MUS | 551 | 551 | 0.683389 | + |
| MUS | 553 | 553 | 0.925689 | + |
| MUS | 556 | 556 | 0.931487 | + |
| MUS | 558 | 558 | 0.691547 | + |
| MUS | 559 | 559 | 0.971129 | + |
| MUS | 560 | 560 | 0.938113 | + |
| MUS | 563 | 563 | 0.954235 | + |
| MUS | 566 | 566 | 0.967467 | + |
| MUS | 567 | 567 | 0.932682 | + |
| MUS | 570 | 570 | 0.951799 | + |
| MUS | 572 | 572 | 0.839915 | + |
| MUS | 573 | 573 | 0.963663 | + |
| MUS | 574 | 574 | 0.928422 | + |
| MUS | 577 | 577 | 0.899768 | + |
| MUS | 579 | 579 | 0.767341 | + |
| MUS | 580 | 580 | 0.946279 | + |

| | | | | |
|---|---|---|---|---|
| MUS | 581 | 581 | 0.89457 | + |
| MUS | 584 | 584 | 0.886062 | + |
| MUS | 586 | 586 | 0.83614 | + |
| MUS | 587 | 587 | 0.939191 | + |
| MUS | 588 | 588 | 0.905539 | + |
| MUS | 591 | 591 | 0.880483 | + |
| MUS | 593 | 593 | 0.856288 | + |
| MUS | 594 | 594 | 0.959942 | + |
| MUS | 595 | 595 | 0.898068 | + |
| MUS | 598 | 598 | 0.773555 | + |
| MUS | 600 | 600 | 0.838638 | + |
| MUS | 601 | 601 | 0.945883 | + |
| MUS | 602 | 602 | 0.902728 | + |
| MUS | 605 | 605 | 0.900001 | + |
| MUS | 607 | 607 | 0.877611 | + |
| MUS | 608 | 608 | 0.962159 | + |
| MUS | 612 | 612 | 0.909255 | + |
| MUS | 614 | 614 | 0.699823 | + |
| MUS | 615 | 615 | 0.909116 | + |
| MUS | 619 | 619 | 0.763155 | + |
| MUS | 624 | 624 | 0.836727 | + |
| MUS | 631 | 631 | 0.957254 | + |
| MUS | 632 | 632 | 0.5 | + |
| MUS | 634 | 634 | 0.560477 | + |
| MUS | 639 | 639 | 0.867656 | + |
| MUS | 641 | 641 | 0.831486 | + |
| MUS | 643 | 643 | 0.864738 | + |
| MUS | 647 | 647 | 0.510238 | + |
| MUS | 648 | 648 | 0.532835 | + |
| MUS | 656 | 656 | 0.70976 | + |
| MUS | 658 | 658 | 0.725052 | + |
| MUS | 662 | 662 | 0.732086 | + |
| MUS | 663 | 663 | 0.682057 | + |
| MUS | 667 | 667 | 0.400653 | |
| MUS | 674 | 674 | 0.731579 | + |
| MUS | 676 | 676 | 0.683573 | + |
| MUS | 681 | 681 | 0.684517 | + |
| MUS | 683 | 683 | 0.602425 | + |
| MUS | 688 | 688 | 0.565729 | + |
| MUS | 690 | 690 | 0.315646 | |
| MUS | 695 | 695 | 0.476853 | |
| MUS | 702 | 702 | 0.0864334 | |
| MUS | 704 | 704 | 0.115475 | |

**Table S8**. Rates of sperm penetration and binding to bovine ZP after homologous (bovine sperm) or heterologous (human and murine sperm) EZPT using bovine ZPs co-incubated without bovine OVGP1 (HoA, HeAh, HeAm) or with bovine OVGP1 (HoB, HeBh, HeBm)

| Group<br>N | Sperm | Bovine OVGP1 | TMC | Penetrated ZPs (%) | Sperm penetration inside ZP (mean) | Max. no. of sperm inside a single ZP | ZP with bound sperm (%) | Sperm bound to ZPs (mean) | Max. no. of sperm bound to a single ZP |
|---|---|---|---|---|---|---|---|---|---|
| HoA<br>67 | Bovine | - | 4.15 ± 0.61 | 67<br>(100 ± 0)[a] | 730<br>(10.89 ± 0.40)[a] | 33 ± 5.59 | 67<br>(100 ± 0) | 918<br>(13.70 ± 0.21)[a] | 25 ± 2.45 |
| HeAh<br>30 | Human | - | 8,81 ± 0.74 | 21<br>(70.0 ± 10.0)[b] | 182<br>(8.66 ± 0.68)[b] | 13 ± 2.08 | 10<br>(33.33 ± 5.77) | 18<br>(0.60 ± 0.66)[b] | 3 ± 0.58 |
| HeAm<br>83 | Murine | - | 8.76 ± 1.07 | 80<br>(96.25 ± 5.3)[c] | 300<br>(3.75 ± 0.31)[c] | 11 ± 2.12 | 83<br>(100 ± 0) | 490<br>(5.90 ± 0.81)[c] | 20 ±3.54 |
| HoB<br>70 | Bovine | + | 4.15 ± 0.61 | 69<br>(98.57 ± 1.36)[a] | 708<br>(10.26 ± 1.51)[a] | 29 ± 4.30 | 70<br>(100 ± 0) | 768<br>(10.97 ± 1.31)[a] | 22 ± 2.38 |
| HeBh<br>90 | Human | + | 8,81 ± 0.74 | 0<br>(0 ± 0)[d] | 0<br>(0± 0)[d] | 0 ± 0 | 12<br>(13.33 ± 3.33) | 12<br>(0.13 ± 0.03)[d] | 1 ± 0 |
| HeBm<br>81 | Murine | + | 8.76 ± 1.07 | 3<br>(3.79 ± 2.08)[d] | 3<br>(1 ± 0)[d] | 1 ± 0 | 36<br>(44.31 ± 3.16) | 94<br>(1.16 ± 0.28)[d] | 6 ± 0.71 |

HoA = homologous EZPT with bovine sperm. HeAh = heterologous EZPT with human sperm. HeAm = heterologous EZPT with murine sperm. HoB = homologous EZPT with bovine sperm (ZPs were co-incubated with bovine OVGP1 for 30 minutes). HeBh = heterologous EZPT with human sperm (ZPs were co-incubated with bovine OVGP1 for 30 minutes). HeBm = heterologous EZPT with murine sperm (ZPs were co incubated with bovine OVGP1 for 30 minutes). TMC = total motile sperm count after capacitation calculated by multiplying the volume by concentration (million sperm/mL) and by the motility (%) after capacitation. Penetrated ZPs refers to the mean number of ZPs penetrated by at least one sperm. ZPs with bound sperm was calculated as the mean number of ZP with, at least, one bound sperm. Sperm penetration was taken as the total number of sperm observed inside the zonas. Bound sperm was expressed as the mean number of sperm that remained bound to the ZP. “N” refers to the total number of ZPs fertilized per treatment. Analysis was conducted by using a one-way ANOVA, followed by Tukey´s post hoc test. Different superscripts (a, b, c, d) in the same column indicate significant differences ($p < 0.05$; n=3). Values are expressed as the mean ± SD.

**Table S9**. Rates of sperm penetration and binding to bovine ZP after homologous (bovine sperm) and heterologous (murine sperm) EZPT using bovine ZPs co-incubated without OVGP1 (HoA, HeA) or with bovine OVGP1 (HoB, HeB). Sperm was also previously co-incubated without OVGP1 or with bovine OVGP1 (HoAb*, HeAb*, HoBb*, HeBb*) or with murine OVGP1 (HeBm*).

| Group<br>N | Sperm | ZPs OVGP1 | Sperm OVGP1 | TMC | Penetrated ZPs (%) | Sperm penetration inside ZP (mean) | Max. no. of sperm inside a single ZP | ZP with bound sperm (%) | Sperm bound to ZPs (mean) | Max. no. of sperm bound to a single ZP |
|---|---|---|---|---|---|---|---|---|---|---|
| HoA<br>15 | Bovine | - | - | 5.74 ± 1.64 | 15<br>(100 ± 0)[a] | 272<br>(18.13 ± 1.32)[a] | 32 ± 2.08 | 15<br>(100 ± 0) | 338<br>(22.53 ± 2.08)[a] | 25 ± 2.51 |
| HoAb*<br>61 | Bovine | - | Bovine | 5.74 ± 1.64 | 61<br>(100.0 ± 10.00)[a] | 963<br>(15.80± 1.18)[a] | 31 ± 1.53 | 61<br>(100 ± 0) | 1093<br>(17.97 ± 3.01)[a] | 31 ± 1.15 |
| HeA<br>60 | Murine | - | - | 4.31 ± 0.94 | 55<br>(91.67 ± 8.15)[b] | 236<br>(4.24 ± 0.52)[b] | 16 ± 1.15 | 60<br>(100 ± 0) | 338<br>(5.63 ± 0.48)[b] | 21 ±1.00 |
| HeAb*<br>55 | Murine | - | Bovine | 4.31 ± 0.94 | 49<br>(88.24 ± 11.76)[b] | 198<br>(4.11 ± 0.51)[b] | 14 ± 1.00 | 55<br>(100 ± 0) | 276<br>(5.00 ± 0.31)[b] | 18 ± 2.65 |
| HeAm*<br>58 | Murine | - | Murine | 4.31 ± 0.94 | 52<br>(89.65 ± 9.75)[b] | 233<br>(4.02 ± 0.66)[b] | 15 ± 1.73 | 58<br>(100 ± 0) | 294<br>(5.08 ± 0.32)[b] | 22 ± 0.58 |
| HoB<br>14 | Bovine | Bovine | - | 5.74 ± 1.64 | 14<br>(100 ± 0)[b] | 232<br>(16.65 ± 0.99)[a] | 33 ± 1.52 | 14<br>(100 ± 0) | 288<br>(20.88 ± 3.88)[a] | 33 ± 2.08 |
| HoBb*<br>59 | Bovine | Bovine | Bovine | 5.74 ± 1.64 | 59<br>(100 ± 0)[a] | 906<br>(15.34 ± 0.66)[a] | 29 ± 1.00 | 59<br>(100 ± 0) | 1001<br>(16.87 ± 2.34)[a] | 32 ± 1.52 |
| HeB<br>16 | Murine | Bovine | - | 4.31 ± 0.94 | 0<br>(0 ± 0)[c] | 0<br>(0 ± 0)[c] | 0 ± 0 | 15<br>(94.44 ± 9.62) | 78<br>(4.90 ± 0.61)[b] | 8 ± 0.57 |
| HeBb*<br>48 | Murine | Bovine | Bovine | 4.31 ± 0.94 | 0<br>(0 ± 0)[c] | 0<br>(0 ± 0)[c] | 0 ± 0 | 41<br>(84.81 ± 10.67) | 115<br>(2,39 ± 0.16)[c] | 11 ± 1.52 |
| HeBm*<br>51 | Murine | Bovine | Murine | 4.31 ± 0.94 | 2<br>(3.98 ± 3.51)[c] | 2<br>(1 ± 0)[c] | 1 ± 0 | 51<br>(100 ± 0) | 229<br>(4.48 ± 0.57)[b] | 9 ± 1 |

HoA = homologous EZPT with bovine sperm. HoAb* = homologous EZPT with bovine sperm (sperm was previously co-incubated with bovine OVGP1 for 30 minutes). HeA = heterologous EZPT with murine sperm. HeAb* = heterologous EZPT with murine sperm (sperm was previously co-incubated with bovine OVGP1 for 30 minutes). HeAm* = heterologous EZPT with murine sperm (sperm was previously co-incubated with murine OVGP1 for 30 minutes). HoB = homologous EZPT with bovine sperm (ZPs were co-incubated with bovine OVGP1 for 30 minutes). HoBb* = homologous EZPT with bovine sperm (ZPs were co-incubated with bovine OVGP1 for 30 minutes) (sperm was previously co-incubated with bovine OVGP1 for 30 minutes). HeB = heterologous EZPT with murine sperm (ZPs were co-incubated with bovine OVGP1 for 30 minutes). HeBb* = heterologous EZPT with murine sperm (ZPs were co-incubated with bovine OVGP1 for 30 minutes) (sperm was previously co-incubated with bovine OVGP1 for 30 minutes). HeBm* = heterologous EZPT with murine sperm (ZPs were co-incubated with bovine OVGP1 for 30 minutes) (sperm was previously co-incubated with murine OVGP1 for 30 minutes). TMC =

total motile sperm count after capacitation calculated by multiplying the volume by concentration (million sperm/mL) and by the motility (%) after capacitation. Penetrated ZP refers to the mean number of ZPs penetrated by at least one sperm. ZPs with bound sperm was calculated as the mean number of ZP with, at least, one bound sperm. Sperm penetration was taken as the total number of sperm observed inside the zonas. Bound sperm was expressed as the mean number of sperm that remained bound to the ZP. “N” refers to the total number of ZPs fertilized per treatment. Analysis was conducted by using a one-way ANOVA, followed by Tukey´s post hoc test. Different superscripts (a, b, c) in the same column indicate significant differences ($p < 0.05$; n=3). Values are expressed as the mean ± SD.

**Table S10**. Rates of sperm penetration and binding to bovine ZP after homologous (bovine sperm) and heterologous (human and murine sperm) EZPT using bovine ZPs co-incubated without OVGP1 (HoA, HeAh, HeAm) or with bovine OVGP1 (HoBb, HeBh, HeBm), human OVGP1 (HoC, HeCh, HeCm) or murine OVGP1 (HoD, HeDh, HeDm)

| Group N | Sperm | OVGP1 | TMC | Penetrated ZPs (%) | Sperm penetration inside ZP (mean) | Max. no. of sperm inside a single ZP | ZP with bound sperm (%) | Sperm bound to ZPs (mean) | Max. no. of sperm bound to a single ZP |
|---|---|---|---|---|---|---|---|---|---|
| HoA 67 | Bovine | - | 4.15 ± 0.61 | 67 (100 ± 0)[a] | 730 (10.89 ± 0.40)[a] | 33 ± 5.59 | 67 (100 ± 0) | 918 (13.70 ± 0.21)[a] | 25 ± 2.51 |
| HeAh 30 | Human | - | 8,81 ± 0.74 | 21 (70.0 ± 10.0)[c] | 182 (8.66 ± 0.68)[c] | 13 ± 2.08 | 10 (33.33 ± 5.77) | 18 (0.60 ± 0.66)[b] | 3 ± 0.58 |
| HeAm 83 | Murine | - | 8.76 ± 1.07 | 80 (96.25 ± 5.3)[a] | 300 (3.75 ± 0.31)[c] | 21 ± 2.12 | 83 (100 ± 0) | 490 (5.90 ± 0.81)[c] | 20 ±3.54 |
| HoB 70 | Bovine | Bovine | 4.15 ± 0.61 | 69 (98.57 ± 1.36)[a] | 708 (10.26 ± 1.51)[a] | 29 ± 4.30 | 70 (100 ± 0) | 768 (10.97 ± 1.31)[a] | 22 ± 2.38 |
| HeBh 90 | Human | Bovine | 8,81 ± 0.74 | 0 (0 ± 0)[b] | 0 (0± 0)[b] | 0 ± 0 | 12 (13.33 ± 3.33) | 12 (0.13 ± 0.03)[b] | 1 ± 0 |
| HeBm 81 | Murine | Bovine | 8.76 ± 1.07 | 3 (3.79 ± 2.08)[b] | 3 (1 ± 0)[b] | 1 ± 0 | 36 (44.31 ± 3.16) | 94 (1.17 ± 0.28)[b] | 6 ± 0.71 |
| HoC 84 | Bovine | Human | 5.03 ± 0.66 | 49 (58.3 ± 1.12)[c] | 235 (4.80 ± 0.02)[c] | 11 ± 1.41 | 84 (100 ± 0) | 820 (4.80 ± 0.02)[c] | 27 ± 0.71 |
| HeCh 87 | Human | Human | 4.79 ± 0.15 | 53 (61.11 ± 7.85)[c] | 249 (4.73 ± 0.73)[c] | 8 ± 0.71 | 87 (100 ± 0) | 832 (9.56 ± 0.02)[c] | 27 ± 1.41 |
| HeCm 30 | Murine | Human | 6.90 ± 0.67 | 0 (0 ± 0)[b] | 0 (0 ± 0)[b] | 0 ± 0 | 2 (6.67 ± 0) | 2 (0.08 ± 0.03)[b] | 1 ± 0 |
| HoD 68 | Bovine | Murine | 4.35 ± 0.46 | 43 (63.16 ± 3.61)[c] | 199 (4.66 ± 0.55)[c] | 7 ± 0.71 | 68 (100 ± 0) | 602 (8.86 ± 0.20)[c] | 16 ± 0.71 |
| HeDh 23 | Human | Murine | 4.79 ± 0.15 | 0 (0 ± 0)[b] | 0 (0± 0)[b] | 0 ± 0 | 1 (12.5 ± 8.84) | 1 (0.06 ± 0.09)[b] | 1 ± 0.71 |
| HeDm 73 | Murine | Murine | 5.23 ± 0.43 | 46 (63.23 ± 7.55)[c] | 211 (4.59 ± 0.13)[c] | 7 ± 1.41 | 73 (100 ± 0) | 980 (13.42 ± 0.27)[a] | 25 ± 3.53 |

HoA = homologous EZPT with bovine sperm. HeAh = heterologous EZPT with human sperm. HeAm = heterologous EZPT with murine sperm. HoB = homologous EZPT with bovine sperm (ZPs were co-incubated with bovine OVGP1 for 30 minutes). HeBh = heterologous EZPT with human sperm (ZPs were co-incubated with bovine OVGP1 for 30 minutes). HeBm = heterologous EZPT with murine sperm (ZPs were co-incubated with bovine OVGP1 for 30 minutes). HoC = homologous EZPT with bovine sperm (ZPs were co-incubated with human OVGP1 for 30 minutes). HeCh = heterologous EZPT with human sperm (ZPs were co-incubated with human OVGP1). HeCm = heterologous

EZPT with murine sperm (ZPs were co-incubated with human OVGP1 for 30 minutes). HoD = homologous EZPT with bovine sperm (ZPs were co-incubated with murine OVGP1 for 30 minutes). HeDh = heterologous EZPT with human sperm (ZPs were co-incubated with murine OVGP1). HeDm = heterologous EZPT with murine sperm (ZPs were co-incubated with murine OVGP1 for 30 minutes). TMC = total motile sperm count after capacitation calculated by multiplying the volume by concentration (million sperm/mL) and by the motility (%) after capacitation. Penetrated ZP refers to the mean number of ZPs penetrated by at least one sperm. ZPs with bound sperm was calculated as the mean number of ZP with, at least, one bound sperm. Sperm penetration was taken as the total number of sperm observed inside the zonas. Bound sperm was expressed as the mean number of sperm that remained bound to the ZP. "N" refers to the total number of ZPs fertilized per treatment. Analysis was conducted by using a one-way ANOVA, followed by Tukey´s post hoc test. Different superscripts (a, b, c) in the same column indicate significant differences ($p < 0.05$; n=3). Values are expressed as the mean ± SD.

**Table S11**. Rates of sperm penetration and binding to murine ZP after homologous (murine sperm) and heterologous (human and bovine sperm) EZPT using bovine ZPs co-incubated without OVGP1 (HoA, HeAh, HeAb) or with in vivo-matured murine OVGP1 (oocytes from superovulated females) (HoBb, HeBh, HeBb), murine OVGP1 (HoC, HeCh, HeCb), human OVGP1 (HoD, HeDh, HeDb) or bovine OVGP1 (HoE, HeEh, HeEb)

| Group N | Sperm | OVGP1 | TMC | Penetrated ZPs (%) | Sperm penetration inside ZP (mean) | Max. no. of sperm inside a single ZP | ZP with bound sperm (%) | Sperm bound to ZPs (mean) | Max. no. of sperm bound to a single ZP |
|---|---|---|---|---|---|---|---|---|---|
| HoA 56 | Murine | - | 8.76 ± 1.08 | 56 (100 ± 0)[b] | 848 (15.14 ± 1.19)[b] | 29 ± 5.50 | 56 (100 ± 0) | 1042 (18.60 ± 2.21) | 32 ± 2.06 |
| HeAh 85 | Human | - | 6.12 ± 0.17 | 60 (70.83 ± 5.89)[c] | 177 (2.95 ± 0.07)[c] | 9 ± 0.71 | 79 (92.92 ± 0.59) | 263 (3.32 ± 0.31)[d] | 11 ± 1.41 |
| HeAb 97 | Bovine | - | 3.67 ± 0.19 | 56 (57.85 ± 5.45)[c] | 167 (2.95 ± 0.02)[c] | 6 ± 0.00 | 91 (93.87 ± 2.65) | 303 (3.32 ± 0.40)[d] | 10 ± 1.41 |
| HoB 19 | Murine | Murine | 5.35 ± 0.07 | 19 (100 ± 0)[b] | 228 (12.00 ± 0.15)[b] | 31 ± 7.78 | 19 (100 ± 0) | 313 (16.47 ± 0.64)[b] | 34 ± 2.83 |
| HeBh 30 | Human | Murine | 6.12 ± 0.17 | 1 (6.67 ± 4.71)[a] | 1 (1 ± 0)[a] | 1 ± 0 | 3 (10.00 ± 4.71) | 3 (1.00 ± 0.00)[a] | 1 ± 0 |
| HeBb 27 | Bovine | Murine | 4.26 ± 0.09 | 1 (6.25 ± 4.42)[a] | 1 (1 ± 0)[a] | 1 ± 0 | 3 (10.80 ± 2.41) | 3 (1.00 ± 0.00)[a] | 1 ± 0 |
| HoC 99 | Murine | Human | 6.33 ± 1.46 | 67 (67.49 ± 2.39)[c] | 228 (3.40 ± 0.24)[c] | 11 ± 1.41 | 97 (98.00 ± 2.82) | 449 (4.63 ± 0.09)[d] | 19 ± 0.71 |
| HeCh 104 | Human | Human | 5.60 ± 0.91 | 67 (64.39 ± 2.32)[c] | 185 (2.76 ± 0.02)[c] | 8 ± 0.71 | 101 (10.38 ± 0.54) | 1079 (10.68 ± 0.54)[c] | 29 ± 2.12 |
| HeCb 30 | Bovine | Human | 6.09 ± 1.31 | 0 (0 ± 0)[a] | 0 (0 ± 0)[a] | 0 ± 0 | 2 (6.67 ± 0) | 2 (1.00 ± 0)[a] | 1 ± 0 |
| HoD 88 | Murine | Bovine | 5.29 ± 0.43 | 56 (63.80 ± 1.49)[c] | 219 (3.91 ± 0.21)[c] | 8 ± 0.71 | 87 (99.01 ± 1.39) | 459 (5.28 ± 0.62)[d] | 19 ± 1.41 |
| HeDh 25 | Human | Bovine | 5.45 ± 0.75 | 0 (0 ± 0)[a] | 0 (0± 0)[a] | 0 ± 0 | 1 (3.33 ± 4.71) | 1 (1.00 ± 0.04)[a] | 1 ± 0.71 |
| HeDb 90 | Bovine | Bovine | 4.35 ± 0.46 | 54 (59.88 ± 1.90)[c] | 182 (3.37 ± 1.38)[c] | 7 ± 0.71 | 90 (100 ± 0) | 956 (10.62 ± 0.08)[c] | 28 ± 1.41 |

HoA = homologous EZPT with murine sperm. HeAh = heterologous EZPT with human sperm. HeAb = heterologous EZPT with bovine sperm. HoB = homologous EZPT with murine sperm (ZPs were co-incubated with murine OVGP1 for 30 minutes). HeBh = heterologous EZPT with human sperm (ZPs were co-incubated with murine OVGP1 for 30 minutes). HeBb = heterologous EZPT with bovine sperm (ZPs were co-incubated with murine OVGP1 for 30 minutes). HoC= homologous EZPT with murine sperm (ZPs were co-incubated with human OVGP1 for 30 minutes). HeCh = heterologous EZPT with human sperm (ZPs were co-incubated with human OVGP1). HeCb = heterologous EZPT with bovine sperm (ZPs were co- incubated with human OVGP1 for 30 minutes). HoD = homologous EZPT with murine sperm (ZPs were co-incubated with bovine OVGP1 for 30 minutes). HeDh = heterologous EZPT with human sperm (ZPs were co-incubated with bovine OVGP1). HeDb = heterologous EZPT with bovine sperm (ZPs were co incubated with bovine OVGP1 for 30 minutes). TMC = total motile sperm count after capacitation calculated by multiplying the volume by concentration (million sperm/mL) and by the motility (%) after capacitation. Penetrated ZP refers to the mean number of ZPs penetrated by at least one sperm. ZPs with bound sperm was calculated as the mean number of ZP with, at least, one bound sperm. Sperm penetration was taken as the total number of sperm observed inside the zonas. Bound sperm was expressed as the mean number of sperm that remained bound to the ZP. "N" refers to the total number of ZPs fertilized per treatment. Analysis was conducted by using a one-way ANOVA, followed by Tukey´s post hoc test. Different superscripts (a, b, c. d) in the same column indicate significant differences ($p < 0.05$; n=3). Values are expressed as the mean ± SD.

**Table S12**. Rates of sperm penetration and binding to bovine ZP after homologous (bovine sperm) EZPT using bovine ZPs co-incubated with neuraminidase (NMase) and without NMase and with bovine OVGP1 or without bovine OVGP1

| Group<br>N | NMase | Bovine OVGP1 | TMC | Penetrated ZPs (%) | Sperm penetration inside ZP (mean) | Max. no. of sperm inside a single ZP | ZP with bound sperm (%) | Sperm bound to ZPs (mean) | Max. no. of sperm bound to a single ZP |
|---|---|---|---|---|---|---|---|---|---|
| HoA<br>55 | + | - | 5.89 ± 0.22 | 2<br>(3.67 ± 0.47)[b] | 2<br>(1 ± 0.0)[b] | 1 ± 0 | 53<br>(96.67 ± 4.71) | 159<br>(2.89 ± 0.35)[b] | 4 ± 0.00 |
| HoAb<br>53 | + | + | 5.89 ± 0.22 | 3<br>(5.89 ± 3.45)[b] | 3<br>(1 ± 0)[b] | 1 ± 0 | 52<br>(98.27 ± 2.44) | 202<br>(3.81 ± 0.73)[b] | 8 ± 0.71 |
| HoB<br>46 | - | - | 5.89 ± 0.22 | 46<br>(100 ± 0)[a] | 492<br>(10.62 ± 0.82)[a] | 33 ± 2.12 | 46<br>(100 ± 0) | 759<br>(16.50 ± 2.56)[a] | 33 ± 2.82 |
| HoBb<br>45 | - | + | 5.89 ± 0.22 | 45<br>(100 ± 0)[a] | 499<br>(11.00 ± 0.82)[a] | 32 ± 1.41 | 45<br>(100 ± 0) | 757<br>(16.82 ± 2.34)[a] | 31 ± 0.71 |

HoA = homologous EZPT with bovine sperm (ZPs were co-incubated with NMase at 10U/mL in acetate buffer, PH 4.5 at 38ºC for 18 hours). HoAb = homologous EZPT with bovine sperm (ZPs were co-incubated with bovine OVGP1 for 30 minutes and then co-incubated with NMase diluted to 10µ/mL in acetate buffer, PH 4.5 at 38ºC for 18 hours). HoB = homologous EZPT with bovine sperm (ZPs were co-incubated with acetate buffer, PH 4.5 at 38ºC for 18 hours). HoBb = homologous EZPT with bovine sperm (ZPs were co-incubated with bovine OVGP1 for 30 minutes and then co-incubated with acetate buffer, PH 4.5 at 38ºC for 18 hours). TMC = total motile sperm count after capacitation calculated by multiplying the volume by concentration (million sperm/mL) and by the motility (%) after capacitation. Penetrated ZPs refers to the mean number of ZPs penetrated by at least one sperm. ZPs with bound sperm was calculated as the mean number of ZP with, at least, one bound sperm. Sperm penetration was taken as the total number of sperm observed inside the zonas. Bound sperm was expressed as the mean number of sperm that remained bound to the ZP. "N" refers to the total number of ZPs fertilized per treatment. Analysis was conducted by using a one-way ANOVA, followed by Tukey´s post hoc test. Different superscripts (a, b) in the same column indicate significant differences ($p < 0.05$; n=3). Values are expressed as the mean ± SD.

**Table S13**. Rates of sperm penetration and binding to murine ZP after homologous (murine sperm) EZPT using murine ZPs co-incubated with neuraminidase (NMase) and without NMase and with murine OVGP1 or without murine OVGP1

| Group N | NMase | Murine OVGP1 | TMC | Penetrated ZPs (%) | Sperm penetration inside ZP (mean) | Max. no. of sperm inside a single ZP | ZP with bound sperm (%) | Sperm bound to ZPs (mean) | Max. no. of sperm bound to a single ZP |
|---|---|---|---|---|---|---|---|---|---|
| HoA 55 | + | - | 6.90 ± 0.94 | 3 (5.45 ± 2.16)[b] | 3 (1 ± 0)[b] | 1 ± 0 | 53 (96.15 ± 5.43) | 257 (4.84 ± 0.59)[b] | 9 ± 0.71 |
| HoAm 53 | + | + | 6.90 ± 0.94 | 3 (5.66 ± 2.31)[b] | 3 (1 ± 0)[b] | 1 ± 0 | 50 (93.98 ± 5.36) | 186 (3.50 ± 1.40)[b] | 10 ± 0 |
| HoB 28 | - | - | 6.90 ± 0.94 | 28 (100 ± 0)[a] | 345 (12.29 ± 0.74)[a] | 33 ± 1.99 | 28 (100 ± 0) | 512 (18.28 ± 0.67)[a] | 31 ± 7.99 |
| HoBm 40 | - | + | 6.90 ± 0.94 | 40 (100 ± 0)[a] | 398 (9,97 ± 0.8)[a] | 31 ± 0,99 | 40 (100 ± 0) | 363 (9.08 ± 0.18)[a] | 30 ± 0.98 |

HoA = homologous EZPT with murine sperm (ZPs were co-incubated with NMase diluted to 10µ/mL in acetate buffer, PH 4.5 at 38ºC for 18 hours). HoAm = homologous EZPT with murine sperm (ZPs were co-incubated with murine OVGP1 for 30 minutes and then co-incubated with NMase at 10U/mL in acetate buffer, PH 4.5 at 38ºC for 18 hours). HoB = homologous EZPT with murine sperm (ZPs were co-incubated with acetate buffer, PH 4.5 at 38ºC for 18 hours). HoBm = homologous EZPT with murine sperm (ZPs were co-incubated with murine OVGP1 for 30 minutes and then co-incubated with acetate buffer, PH 4.5 at 38ºC for 18 hours). TMC = total motile sperm count after capacitation calculated by multiplying the volume by concentration (million sperm/mL) and by the motility (%) after capacitation. Penetrated ZPs refers to the mean number of ZPs penetrated by at least one sperm. ZPs with bound sperm was calculated as the mean number of ZP with, at least, one bound sperm. Sperm penetration was taken as the total number of sperm observed inside the zonas. Bound sperm was expressed as the mean number of sperm that remained bound to the ZP. "N" refers to the total number of ZPs fertilized per treatment. Analysis was conducted by using a one-way ANOVA, followed by Tukey´s post hoc test. Different superscripts (a, b) in the same column indicate significant differences ($p < 0.05$; n=3). Values are expressed as the mean ± SD.

**Table S14**. Rates of sperm penetration and binding to bovine ZP after homologous (bovine sperm) and heterologous (murine sperm) EZPT using bovine ZPs co-incubated with bovine OVGP1 previously treated with Neuraminidase (NMase) (HoA, HeA) and with non treated bOVGP1 (HoB, HeB). Control groups without ZPs treatment was added for both, homologous (HoC) and heterologous (HeC).

| Group<br>N | Bovine OVGP1 | NMase treatment to bOVGP1 | TMC | Penetrated ZPs (%) | Sperm penetration inside ZP (mean) | Max. no. of sperm inside a single ZP | ZP with bound sperm (%) | Sperm bound to ZPs (mean) | Max. no. of sperm bound to a single ZP |
|---|---|---|---|---|---|---|---|---|---|
| HoA<br>15 | + | + | 5.25 ± 0.60 | 76<br>(84.49 ± 9.91)[a] | 473<br>(5.11 ± 1.12)[a] | 22 ± 1.53 | 87<br>(100 ± 0) | 720<br>(8.02 ± 1.06)[a] | 29 ± 3.05 |
| HoB<br>87 | + | - | 5.25 ± 0.60 | 15<br>(100 ± 0)[b] | 254<br>(21.07 ± 7.61)[b] | 29 ± 3.05 | 15<br>(100 ± 0) | 355<br>(23.67 ± 7.11)[b] | 33 ± 5.51 |
| HeA<br>82 | + | + | 5.34 ± 0.05 | 45<br>(52.78 ± 12.73)[c] | 123<br>(1.46 ± 0.27)[c] | 6.00 ± 1.00 | 65<br>(79.58 ± 7.88) | 286<br>(3.51 ± 0.31)[c] | 11 ± 1.52 |
| HeB<br>15 | + | - | 5.34 ± 0.05 | 0<br>(0 ± 0)[d] | 0<br>(0 ± 0)[d] | 0 ± 0 | 4<br>(26.67 ± 11.54) | 5<br>(0.33 ± 0.23)[d] | 2 ± 0.57 |
| HeC<br>20 | - | - | 5.34 ± 0.05 | 18<br>(88.10 ± 6.73)[a] | 113<br>(5.70 ± 0.19)[a] | 13 ± 2.12 | 20<br>(100 ± 0) | 185<br>(9.98 ± 2.61)[a] | 19 ± 3.54 |

HoA = homologous EZPT with bovine sperm (ZPs were co-incubated with bovine OVGP1 for 30 minutes. Bovine OVGP1 was previously treated by incubation with NMase at 10U/mL in acetate buffer, PH 4.5 at 38ºC for 18 hours). HoB = homologous EZPT with bovine sperm (ZPs were co-incubated with bovine OVGP1 for 30 minutes. Bovine OVGP1 was previously incubated in acetate buffer, PH 4.5 at 38ºC for 18 hours). HeA = heterologous EZPT with murine sperm (ZPs were co-incubated with bovine OVGP1 for 30 minutes. Bovine OVGP1 was previously treated with NMase at 10U/mL in acetate buffer, PH 4.5 at 38ºC for 18 hours). HeB = Heterologous EZPT with murine sperm (ZPs were co-incubated with bovine OVGP1 for 30 minutes. Bovine OVGP1 was previously treated in acetate buffer, PH 4.5 at 38ºC for 18 hours). HeC = heterologous EZPT with murine sperm (ZPs were co-incubated with acetate buffer, PH 4.5 at 38ºC for 18 hours). TMC = total motile sperm count after capacitation calculated by multiplying the volume by concentration (million sperm/mL) and by the motility (%) after capacitation. Penetrated ZPs refers to the mean number of ZPs penetrated by at least one sperm. ZPs with bound sperm was calculated as the mean number of ZP with, at least, one bound sperm. Sperm penetration was taken as the total number of sperm observed inside the zonas. Bound sperm was expressed as the mean number of sperm that remained bound to the ZP. "N" refers to the total number of ZPs fertilized per treatment. Analysis was conducted by using a one-way ANOVA, followed by Tukey´s post hoc test. Different superscripts (a, b, c, d) in the same column indicate significant differences ($p < 0.05$; n=3). Values are expressed as the mean ± SD.
